# Identification of regulatory networks associated with anti-HIV/AIDS genes via integrated transcriptome, epigenome and proteome analyses

**DOI:** 10.1101/2021.03.21.436300

**Authors:** Gexin Liu, Chunlin Zhang, Lei Shi, Zhenglin Zhu

**Affiliations:** School of Life Sciences, Chongqing University, Chongqing, China

**Keywords:** HIV-resistant, AIDS, transcriptome, epigenomics

## Abstract

There are individuals naturally resistant to HIV. To identify anti-HIV genes and regulatory networks that enable the native ability to resist HIV, we reanalyzed the transcriptome of HIV resistant and susceptible individuals based on previous efforts, and performed regulatory network prediction using HIV-infection related DNA methylation, miRNA and Chip-SEQ data. We totally found 25 potential anti-HIV genes and 23 of them are newly identified. They are enriched in pathways of immunity, neurological system and cell signaling. 4 anti-HIV genes show DNA hypermethylation signatures and 4 are possibly bounded by the HIV-1 Trans-Activator of Transcription protein (Tat). We found a potential HIV-resistance correlated miRNA hsa-miR-3074-5p possibly regulating an anti-HIV hub gene JUN. Our findings provide novel insights for AIDS treatments and approaches to HIV vaccine design.

## Introduction

The acquired immunodeficiency syndrome, AIDS [1], as an epidemic and pandemic disease, has caused 33 million reported deaths in recent 40 years, after the first identification of HIV, the human immunodeficiency virus [2], Global statistics show that there are 38 million HIV infections, 1.7 million new infections and 690 thousands AIDS-related confirmed deaths in 2019 [3]. HIV/AIDS remains a serious threat to human health. As a retrovirus and a lentivirus, HIV has two subtypes, HIV-1 and HIV-2. Compare to HIV-2, HIV-1 is more infectious and aggressive. HIV targets and destroys human CD4+ T cells [4–6], disrupting the human immune system and rendering it susceptible to exogenous infectious diseases, such as tuberculosis, [7] and tumors [8]. Until now, there is still no effective medicine to radically cure AIDS nor vaccine to prevent HIV infection [9]. A major treatment against AIDS is the highly active antiretroviral therapy (HAART), which jointly uses three or more antivirus medicines to prevent the replication of HIV virus [10]. This therapy can reduce the virus load (VL) in plasma to a small level, usually undetectable by normal HIV detection methods. However, the virus load will bounce from hidden infected cells if the therapy is terminated [11]. HAART cannot radically cure AIDS. Currently developed HIV vaccines, such as RV144 [12], are capable of preventing the infection of HIV to some extent but are far from effective. The obstacles are the fast mutation of HIV populations [13] and the lack of the understanding of the complex interaction between HIV and human immune system.

With the spread of HIV around the world, it is observed that some individuals are naturally resistant to HIV. Some Caucasian populations have a high frequency allele in CCR5, because of which they are reported to have native immunity to HIV [14–16]. Moreover, in Kenya, Africa, some sex workers are found to be naturally resistant to HIV infection [17, 18]. Previous efforts with comparative proteome analysis of the HIV-resistant individuals and HIV-negative controls have already identified several anti-proteases. The expressions of these anti-proteases are relatively high in HIV-resistant individuals [19, 20]. These anti-proteases are also validated to play a role in the process of HIV-infection [19]. In spite of these efforts, the genomic data of HIV-resistant individuals/populations and related analysis have not been publicly reported to date. Understanding the molecular mechanism of resistance to HIV infection in this population under continued exposure should be important for medicine/vaccine development to prevent HIV infection and to cure AIDS.

With the development of next-generation sequencing technology, more HIV/AIDS related big data are generated [21], such as DNA methylation [22] and ChIP-Seq [23] data. Integrated analysis of multiple data may give new results. For example, a re-analysis of gene expression and methylation regulation in glioblastoma identified an eight-gene signature, which is of prognostic value for glioblastoma patients [24], Following previous efforts [25–28], we collected and jointly analyzed HIV related transcriptomic, epigenomic and proteomic data. In this manuscript, we will show newly identified anti-HIV genes, HIV-resistance pathways, regulatory networks and related potential clinical biomarkers.

## Materials and methods

### Microarray and ChIP-Seq data

We downloaded the microarray data (GSE33580, GSE29429, GSE14278, GSE73968), miRNA data (GSE103555), DNA-methylation data (GSE67705) and ChIP-Seq data (GSE30738) from the GEO database of NCBI. GSE33580 includes 43 HIV-resistant and 43 healthy-control Kenya people’s blood samples. GSE29429 includes 30 HIV-infected and 17 healthy African people’s blood samples. The experiment is performed in the platform GPL10558, Illumina HumanHT-12 V4.0 expression beadchip. GSE14278 includes 9 HIV-resistant and 9 normal Kenya people’s CD4+T cell samples. GSE73968 includes 3 HIV-infected patients’ naive CD4+T cell samples, 3 healthy people’s naive CD4+T cell samples, 3 HIV-infected patients’ central memory CD4+T cell samples and 3 healthy patients’ central memory CD4+T cell samples. The experiment is performed in the platform GPL6244, Affymetrix Human Gene 1.0 ST Array. The platform for GSE33580 and GSE14278 are both GPL570, Affymetrix Human Genome U133 Plus 2.0 Array.

For miRNA data, GSE103555 (Platform GPL18058) includes 10 new HIV-infected cases and 10 old HIV-infected cases and 10 healthy controls. GSE67705 is a DNA-methylation data. It includes 142 chronic HIV-infected patients’ and 44 healthy people’s blood samples. It is performed in the platform GPL13534, Illumina HumanMethylation450 BeadChip. GSE30738 is a ChIP-Seq data, showing the genome-wide binding map of HIV-1 Trans-Activator of Transcription (Tat) to the human genome in Jurkat T cells (Jurkat-Tat cells). It includes ChIP on chip for H3K9ac in Jurkat-Tat versus Jurkat cells.

### The processing of transcriptomic microarray data

We used Affy [29] to process the 43 HIV-infected and 43 healthy Kenya people’s blood samples in GSE33580. We discarded samples that has bad sequencing quality. We used GEO2R to call differently expressed genes (DEGs) requiring p<0.05 and | logFC|>1. We also used Affy to check the quality of 9 HIV-resistant and 9 normal Kenya CD4+T cell samples in GSE14278. We used GEO2R to filter out differently expressed genes (DEGs). We looked for the intersection of the DEGs retrieved from GSE33580 and GSE14278. We also used GEO2R to look for DEGs through comparing 30 HIV-infected and 17 normal African people’s blood samples as the control. In the same way, we did quality check and DEG calling for the HIV-infected and normal CD4+T cell sample data in GSE73968. We merged the DEGs resulted from analyses in the naive CD4+T cell and the DEGs resulted from analyses in the central memory CD4+T cell.

### The processing of DNA-methylation and miRNA microarray data

We used GEO2R to find out differently methylated genes (DMGs) requiring p<0.05 and |t|>2 in the DNA-methylation data GSE67705. In combination with the resulted retrieved from GSE33580, we filtered out DEGs that are down-regulated in transcription but up-regulated in DNA methylation. We also filtered out DEGs that are up-regulated in transcription but down-regulated in DNA methylation. We did the same thing for the DEGs both in blood sample (GSE33580) and CD4+ T cells (GSE14278) for hypermethylated down-regulated genes and hypomethylated up-regulated genes.

We used GEO2R to process miRNA data GSE103555 aiming to find differently expressed miRNAs. We set the cutoffs p<0.01 and |logFC|>2. We predicted the target genes of miRNAs using miRDB [30] and compared the results and the DEGs resulted from the analysis in GSE33580. Finally, we generated a list of predicted relationships between differently expressed miRNA and DEGs.

### The process of ChIP-Seq data

We used fastq-dump to transform SRA format data into fastq format and then preprocess these fastq files by TrimGalore (github.com/FelixKrueger/TrimGalore) requiring Phred quality score > 20 and reads length > 20. Thereafter, we used fastqc [31] to do quality control. We used bowtie2 [32] to map reads onto the human genome UCSC GRCh37/hg19 retrieved from Ensembl [33]. Then used MACS2 [34] to detect the enriched region in the ChIP-Seq data. Finally, we used the ChIPseeker package [35] by R to annotate peaks, within a region 3000 bp near the ChIP-Seq enriched region. We selected out the genes that not only located within the peak region but also DEGs resulted from GSE33580 (Blood-HIV-resistant). We did the same thing for the shared DEGs both in blood sample (GSE33580) and CD4+ T cells (GSE14278).

### GO and pathway enrichment analysis

We used the R package clusterProfiler [36] to do GO and KEGG analysis and draw figures. We did analysis for all DEGs, up-regulated DEGs with high DNA-methylation expression, down-regulated DEGs with low DNA-methylation expression and ChIP-Seq correlated genes.

### Protein-protein interaction network and hub gene selection

We used the Search Tool for the Retrieval of Interacting Genes (String) [37] to generate protein-protein interaction networks and used Cytoscape [38] to view the results. We used cytoHubba [39] in Cytoscape to search for hub genes by algorithms MNC, Degree, EPC, BottleNeck, Closeness and Radiality. We selected out cases with high scores (5% in ranking). We also looked for the overlapped DEGs resulted from GSE33580 (EXP-Blood-HIV-resistant) and GSE14278 (EXP-CD4-HIV-resistant).

### Other analysis

We used Haploview [40] to get, view and analyze SNP information of genes. We utilized a NCBI tool HPA RNA-Seq normal tissues to search for the expression information of genes in different tissues. We used the Connective Map (CMAp) to look for potential small molecular drugs based on DEGs. We filtered out results requiring the score<0 and P-value<0.01.

## Results

### Identification of anti-HIV genes

To identify anti-HIV genes, we analyzed differently expressed genes (DEGs) through comparing the gene expression in the blood of HIV-resistant individuals (HIV-R) and HIV-negative controls (HIV-N-C) based on a published microarray dataset (NCBI GEO ID, GSE33580). For convenience, we named the experiment EXP-Blood-HIV-Resistance in the subsequent description. HIV-resistant samples are collected from highly exposed seronegative (HESN) populations [25]. Following previous efforts [25], we performed strict quality control for sequencing data by relative log expression (RLE) plot. RLE can display the consistency of gene expressions. In the RLE plot, the center of each sample should be very close to the position of ordinate 0. An individual sample significantly different from most of the other samples is always problematic and suggested to be discarded [41], We excluded 13 samples with low quality in EXP-Blood-HIV-Resistance (Figure S1). We next performed analysis for the left 73 samples by GEO2R and identified 452 DEGs, including 81 up-regulated, and 371 down-regulated DEGs in HIV-R (Figure S2, Tables S1). We also performed analyses of DEGs between HIV-R and HIV-N-C based on a documented CD4+ T cell experiment (NCBI GEO ID, GSE14278), named EXP-CD4-HIV-Resistance in post description. We discarded one sample in the quality control (Figure S3) and finally identified 553 DEGs, including 279 up-regulated and 274 down-regulated genes in HIV-R (Figure S4, Table S2). The intersection of the DEGs resulted from EXP-Blood-HIV-Resistance and EXP-CD4-HIV-Resistance is 27 (Figure1A, Figure S5), in which 12 show consistent expression patterns in both cohorts (Table S3). We performed a survey of the 12 DEGs in different tissues based on a published gene expression profiling dataset [42] and found brain, kidney, spleen, testis and adrenal are hot tissues for the expression of the 12 potential anti-HIV genes (Figure1B). For convenience, we used ‘the shared 12 anti-HIV DEGs’ to denote the 12 DEGs with similar expression patterns both in EXP-Blood-HIV-Resistance and EXP-CD4-HIV-Resistance in post description.

**Figure 1.**
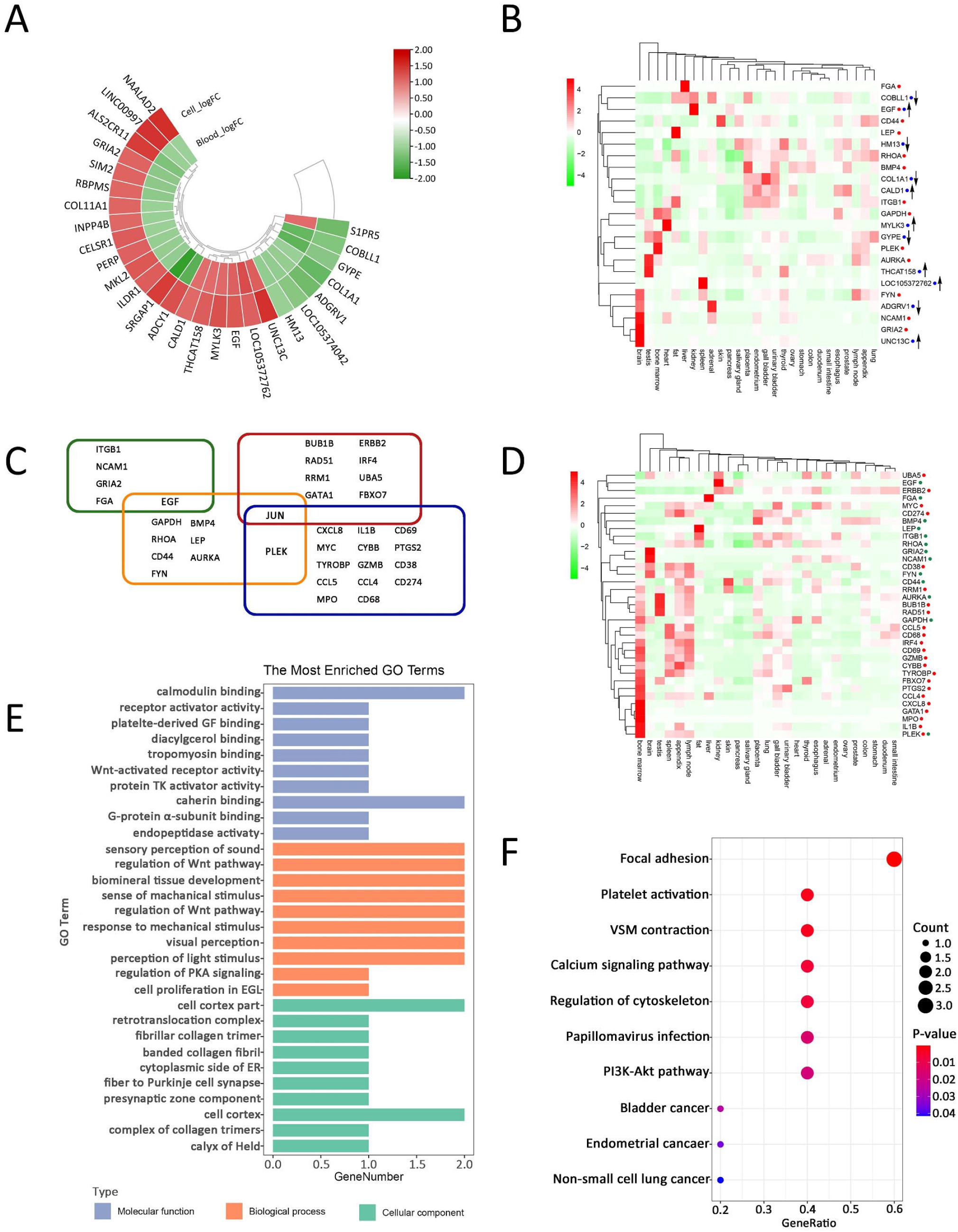
The expression and function of anti-HIV genes. A, a circular heat map showing the expression of the identified anti-HIV genes in EXP-Blood-HIV-Resistance and EXP-CD4-HIV-Resistance. B, a heat map showing the expression in tissues for the identified 25 anti-HIV genes, including the shared 12 anti-HIV DEGs, marked by blue cycles following gene names, and the 14 anti-HIV hub genes, marked by red cycles following gene names. Up and down arrows denote up- and down-regulation, respectively. C, the Venn plot of 36 hub genes in 4 sets of transcriptome data. Those within a green line are hub genes resulted from EXP-Blood-HIV-Resistance, those within an orange line are hub genes resulted from EXP-CD4-HIV-Resistance, those within a red line are hub genes resulted from EXP-Blood-HIV-Infection, and those within a blue line are hub genes resulted from EXP-CD4-HIV-Infection. D, a heat map showing the expression of 36 hub genes in tissues. The hub genes that function in HIV infection are marked by red cycles following gene names, while those function in HIV-resistance are marked by green cycles following gene names. E and F are the plots of GO enrichment analysis and pathway enrichment analysis of the shared 12 anti-HIV DEGs, respectively.

We searched for hub genes by cytoHubba in Cytoscape [43] and predicted the networks of protein-protein interaction of DEGs by STRING [44], We retrieved the top 30 in the score rankings resulted from six approaches, MNC, Degree, EPC, BottleNeck, Radiality and Closeness, provided by cytoHubba [39], and then took the crossing (Table S4). We identified 5 hub genes in the DEGs of EXP-Blood-HIV-Resistance (Figure S6) and 10 hub genes in the DEGs of EXP-CD4-HIV-Resistance (Figure S7). The 14 identified hub DEGs showed high expression in brain, fat, bone marrow, liver, kidney, testis and placenta (Figure1B). Together with the shared 12 anti-HIV DEGs, we in total identified 25 potential anti-HIV genes (Table S5).

For investigating the function of these potential anti-HIV genes in response to HIV exposure, we searched for DEGs between the blood samples of patients with early acute HIV infection (HIV+) and uninfected controls (HIV-) (NCBI GEO ID, GSE29429; named EXP-Blood-HIV-Infection for presentation convenience in following description) and CD4+ T level (NCBI GEO ID, GSE73968; named EXP-CD4-HIV-Infection). We performed quality control for this data (Figure S8, S9). Based on EXP-Blood-HIV-Infection, we identified 1458 DEGs in HIV+ patients, including 925 up-regulated and 533 down-regulated genes (Figure S10, Table S6). Based on EXP-CD4-HIV-Infection, we identified 274 DEGs for naive CD4+ T cell samples, including 192 up-regulated and 82 down-regulated genes (Figure S11), and 152 DEGs for central memory CD4+ T cell samples, including 119 up-regulated and 33 down-regulated genes (Figure S12). In total, there are 358 DEGs with 248 up-regulated and 100 down-regulated genes in EXP-CD4-HIV-Infection (Table S7). There are 43 up-regulated and 4 down-regulated DEGs showing consistent regulation patterns in both experiments (Table S8). Using the same method to search for HIV-R hub DEGs, we searched for HIV+ hub DEGs and identified 9 hub genes in the DEGs of EXP-Blood-HIV-Infection (Figure S13) and 16 hub genes in EXP-CD4-HIV-Infection (Figure S14). Taking together, there are 36 HIV+ hub genes (Figure1C). We constructed the protein-protein interaction (PPI) network of the 36 hub genes (Figure2) and investigated the expression of these genes in different tissues (Figure1D). The results show that JUN and PLEK are two hub genes both in HIV-resistance and HIV-infection. HIV-infection hub genes show biased high expression in the bone-marrow, compared to HIV-resistance hub genes (P-value<0.044, Wilcoxon Test).

**Figure 2.**
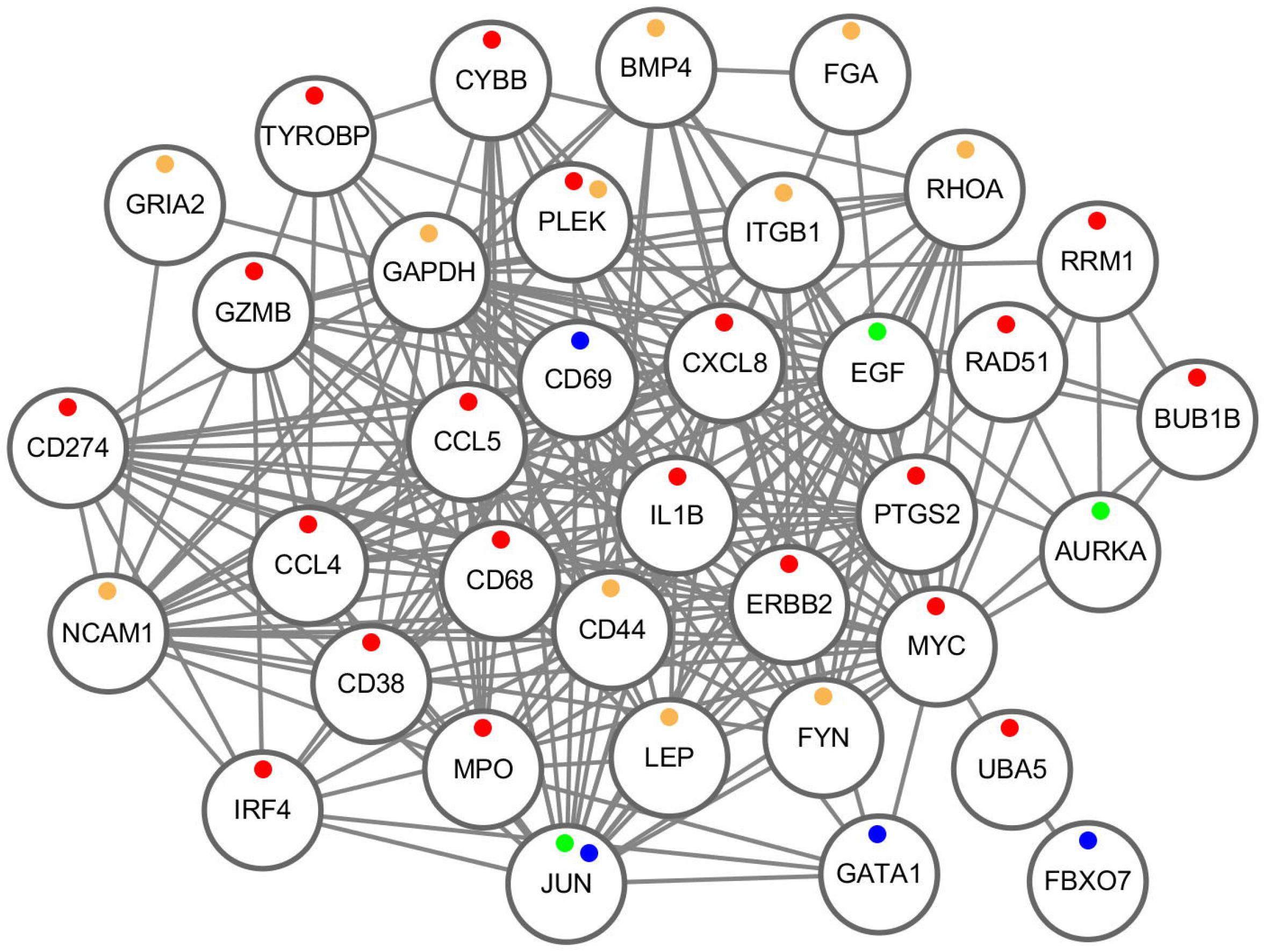
The PPI network diagram of 36 HIV related hub genes. Red dots represent up-regulated HIV-infection genes, blue dots represent down-regulated HIV-infection genes, dark yellow dots represent down-regulated anti-HIV genes and green dots represent up-regulated anti-HIV genes.

### Prediction of anti-HIV regulatory networks

Since the lack of HIV-resistance epigenomic data, we used documented DNA-methylation and miRNA microarray data of HIV-infected samples to predict the potential regulatory network of anti-HIV genes. We searched for differently methylated genes (DMGs) through methylome-wide analysis of chronic HIV infected patients (HIV+), compared with healthy controls (HIV-), using a published data (NCBI GEO ID, GSE67705) and identified 13626 candidates with P-value<0.05 and |t|>2. In the 13626 DMGs, there are 6861 up-regulated genes and 6765 down-regulated genes in HIV+ patients (Table S9). We searched for the overlap between these DMG candidates and the HIV-R DEGs resulted from EXP-Blood-HIV-Resistance. We found 22 up-regulated (HIV-R) and hypomethylated (HIV+) genes (Figure S15, Table S10) and 103 down-regulated (HIV-R) and hypermethylated (HIV+) genes (Figure S16, Table S11). We also searched for the overlap between the DMG candidates and the 12 shared DEGs, and identified 2 up-regulated (HIV-R) and hypomethylated (HIV+) genes but none down-regulated (HIV-R) and hypermethylated (HIV+) gene (Table S12). In addition, we searched for the overlap between the DMG candidates and the 14 HIV-resistant hub genes. We identified one up-regulated (HIV-R) and hypomethylated (HIV+) gene and two down-regulated (HIV-R) and hypermethylated (HIV+) genes (Table S13). In total, there are 125 HIV-R DEGs identified with DNA methylation signatures, in which 103 is hypomethylation and 22 is hypermethylation (Table S10, S11). In all identified anti-HIV genes (Table S5), there are one up-regulated (HIV-R) and hypomethylated (HIV+) gene and 4 down-regulated (HIV-R) and hypermethylated (HIV+) genes (Table S14).

We analyzed differently expressed miRNA (DEM) based on the data between HIV+ and HIV- (NCBI GEO, GSE103555), and identified 47 DEMs, in which 16 are up-regulated and 31 are down-regulated in HIV+ patients (Table S15). Thereafter, we performed target gene prediction by miRDB [30] with the prediction score > 60. We only kept the resulted pairs with the miRNA and the predicted target gene in opposite expression [45]. We tried to identify the regulation patterns of the identified 25 anti-HIV genes using the identified DEMs, and finally obtained a regulatory network including 5 miRNA and 7 anti-HIV genes (Table S16, S17). Hsa-miR-3074-5p is a potential anti-HIV related functional miRNA. Predictions show that hsa-miR-3074-5p may negatively regulate the expression of JUN, an anti-HIV and HIV+ related hub gene shared by 4 cohorts (Figure 1C). The up-regulation of JUN may benefit HIV-resistance (Table S2). Thus, the inhibition of this miRNA may help to resist HIV infection.

We also searched for the genes bounded by the HIV-1 Trans-Activator of Transcription protein (Tat). We performed analysis on a documented ChIP-Seq data (GSE30738), which includes a genome-wide binding map of the HIV Tat protein to the human genome. We identified 5344 Tat binding genes (Table S18), in which 124 are DEGs in EXP-Blood-HIV-Resistance (Table S19), and 4 are DEGs both in EXP-Blood-HIV-Resistance and EXP-CD4-HIV-Resistance (Table S20).

### Functional prediction of anti-HIV genes

We used the R package cluster Profiler to do gene ontology (GO) and pathway enrichment analyses (for details, see Materials and methods). The 452 DEGs in EXP-Blood-HIV-Resistance and the 553 DEGs in EXP-CD4-HIV-Resistance share a lot. They are both enriched in the function of extracellular biological factor binding, channel activity, cell signaling transduction and cell differentiation (Figure S17, S18; Table S21, S22). They are also both enriched in pathways associated with blood circulatory, neurological diseases and cancer (Figure S19, S20; Table S23, S24). DEGs in the blood or the cells extracted from HIV-infected patients also participated in functions or pathways similar to HIV-resistant DEGs (Figure S21-S24, Table S25-S28). The shared 12 anti-HIV DEGs function in focal adhesion, platelet activation and vascular smooth muscle contraction (Figure 1E, F; Table S29, S30). For HIV-R DEGs showing different DNA methylation, down-regulated HIV-R DEGs with hypermethylation show an enrichment in neurological activities (Figure S25, S26; Table S31, S32) while up-regulated HIV-R DEGs with hypomethylation show an enrichment in cell cycle (Figure S27, S28; Table S33, S34). Tat binding HIV-R DEGs in EXP-CD4-HIV-Resistance show a preference to function in cell cycle (Figure S29, S30; Table S35, S36). In pathway enrichment analysis of anti-HIV DEGs, we identified 5 potential anti-HIV pathways (Table S37), including focal adhesion (Figure 3), regulation of actin cytoskeleton (Figure S31), platelet activation (Figure S32), cAMP signaling (Figure S33) and gastric acid secretion (Figure S34).

**Figure 3.**
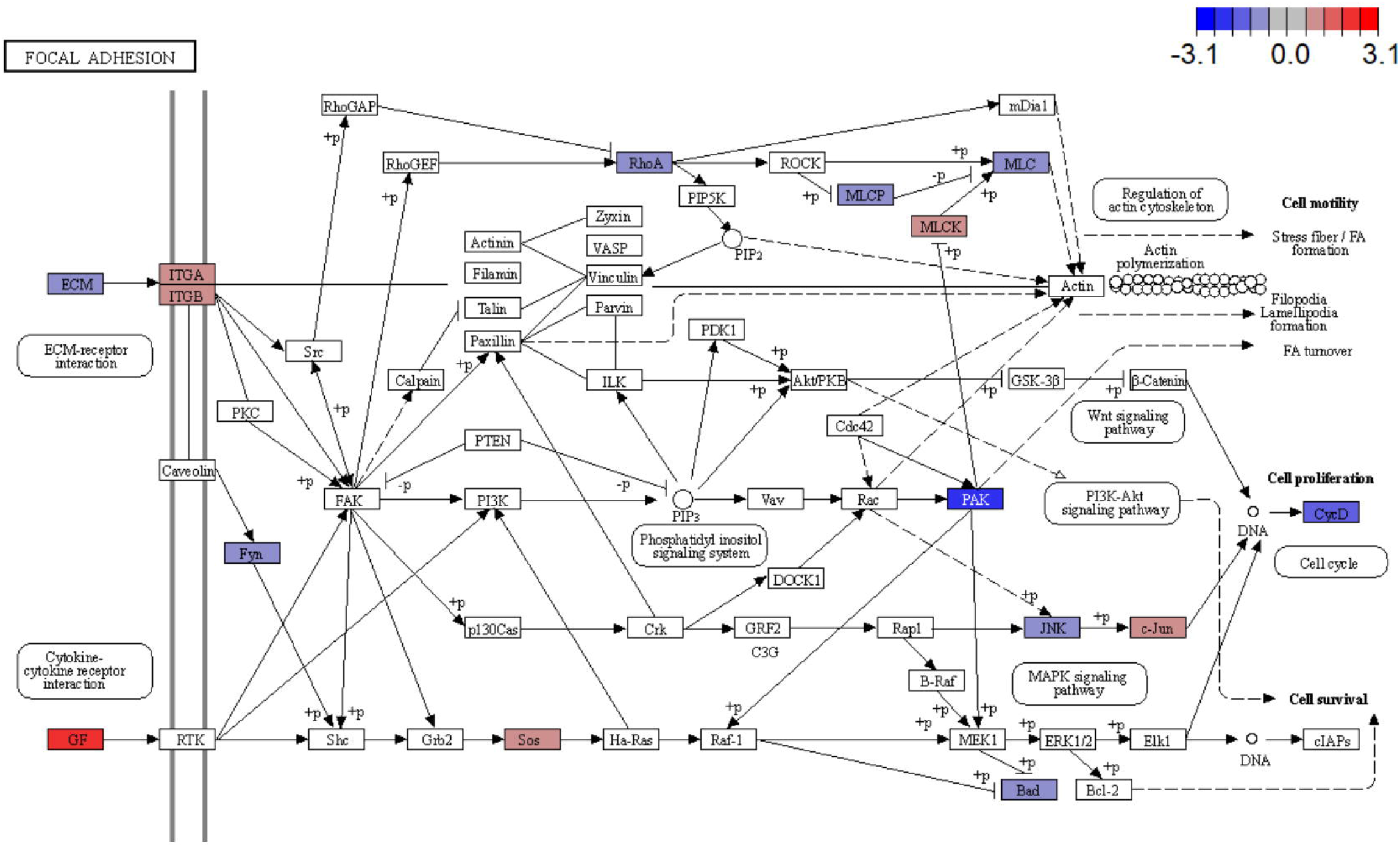
The mapped DEGs in the pathway, focal adhesion. It is resulted from a common pathway enrichment analysis based on EXP-Blood-HIV-Resistance and EXP-CD4-HIV-Resistance. Mapped DEGs are in colored rectangles. A color akin to red denotes up-regulation, while that akin to blue denotes down-regulation.

### Potential anti-HIV biomarkers

JUN and PLEK are the only two hub genes shared by HIV resistance and HIV infection cohorts. There is no previous report of HIV resistance for these two genes. JUN is HIV-R up-regulated but HIV+ down-regulated (Table S2, S6). It can be inferred that the increase in the expression of JUN may promote the resistance to HIV-infection, that individuals with low expression level of JUN may be susceptible to HIV infection, and that HIV-infection may lead to the reduced expression of JUN. The protein encoded by JUN is a major component of the AP-1 complex, which commonly functions in growth and differentiation. It is also reported that the transcription of JUN is stimulated by its own gene products [46, 47], PLEK show low expression level in HIV-R but high in HIV+ (Table S2, S6). The silence of PLEK may benefit resistance in exposure to HIV. PLEK encodes a protein named Pleckstrin, which is the main substrate of the protein kinase C (PKC), highly expressed in platelets and white blood cells [48, 49]. Studies have also provided evidence that supports the role of PLEK in exocytosis [50, 51].

EGF is the only anti-HIV hub gene shared by EXP-Blood-HIV-Resistance and EXP-CD4-HIV-Resistance. This gene is a HIV-R up-regulated DEG in both experiments. We performed a research in population genetics for EGF and identified 75 SNP tags (Table S38). We used the human population genomic data of Kenya. The reason is that the samples in EXP-Blood-HIV-Resistance and EXP-CD4-HIV-Resistance were collected from African people. Through using CMap [52] to analyze the shared 12 anti-HIV DEGs, we found 12 small molecules as potential therapeutic drugs for HIV (Table S39).

## Discussion

We performed a comprehensive analysis of HIV-resistance transcriptomic and epigenomic data (Figure 4) and identified 25 potential anti-HIV genes and constructed a network based on our analyses (Figure 5). There are 12 DEGs with similar regulation patterns (6 up-regulation and 6 down-regulation in HIV-R) both in EXP-Blood-HIV-Resistance and CD4-HIV-Resistance, and 15 DEGs showing pairs of opposite regulation patterns in the two experiments. These may be leaded by the difference between blood sample and CD4+ T cells, or by the sampling error.

**Figure 4.**
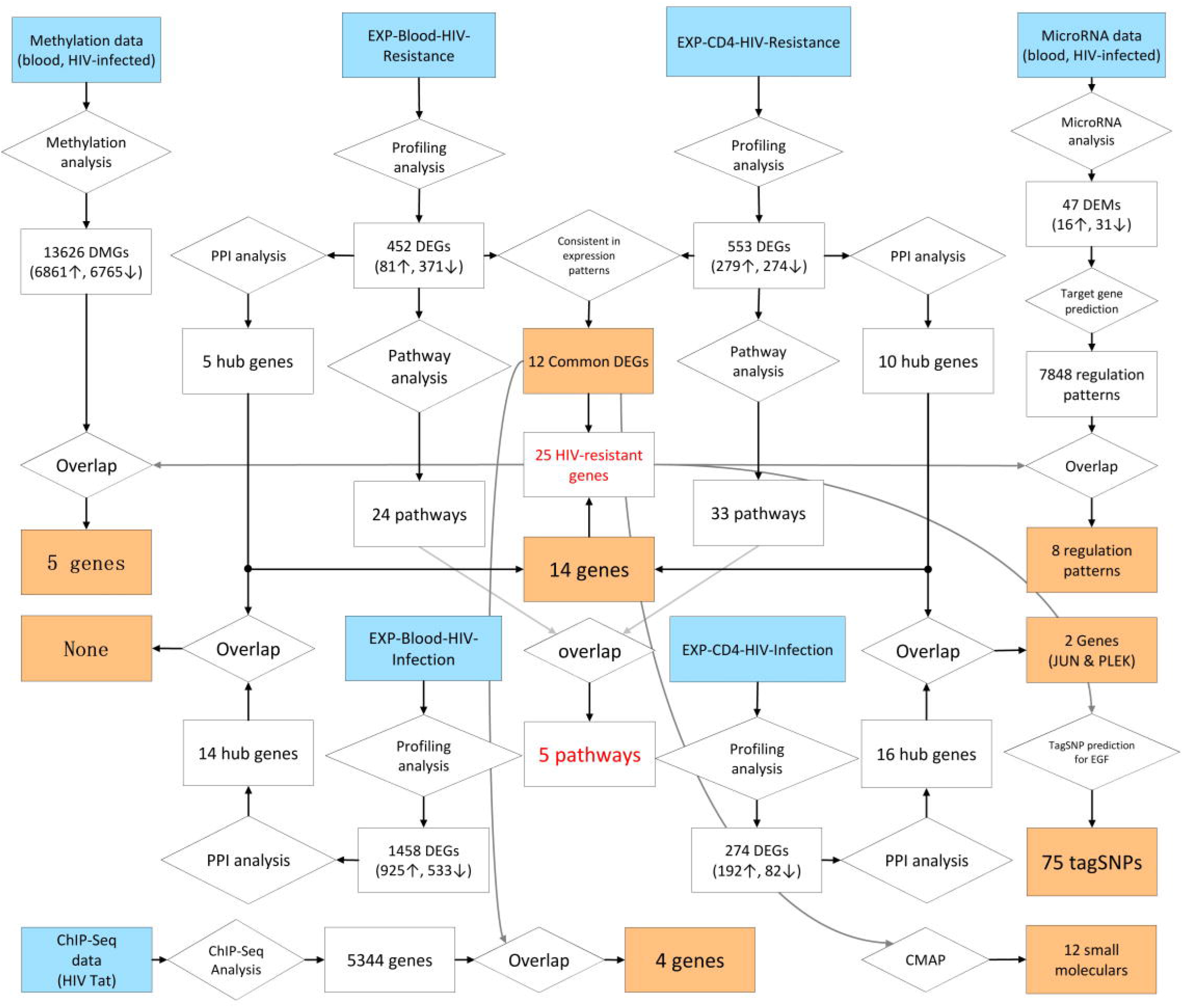
The flow chart of the entire analysis process. Boxes in blue denote the beginning of the pipeline. Boxes in orange denote the results, and diamond boxes denote analysis steps.

**Figure 5.**
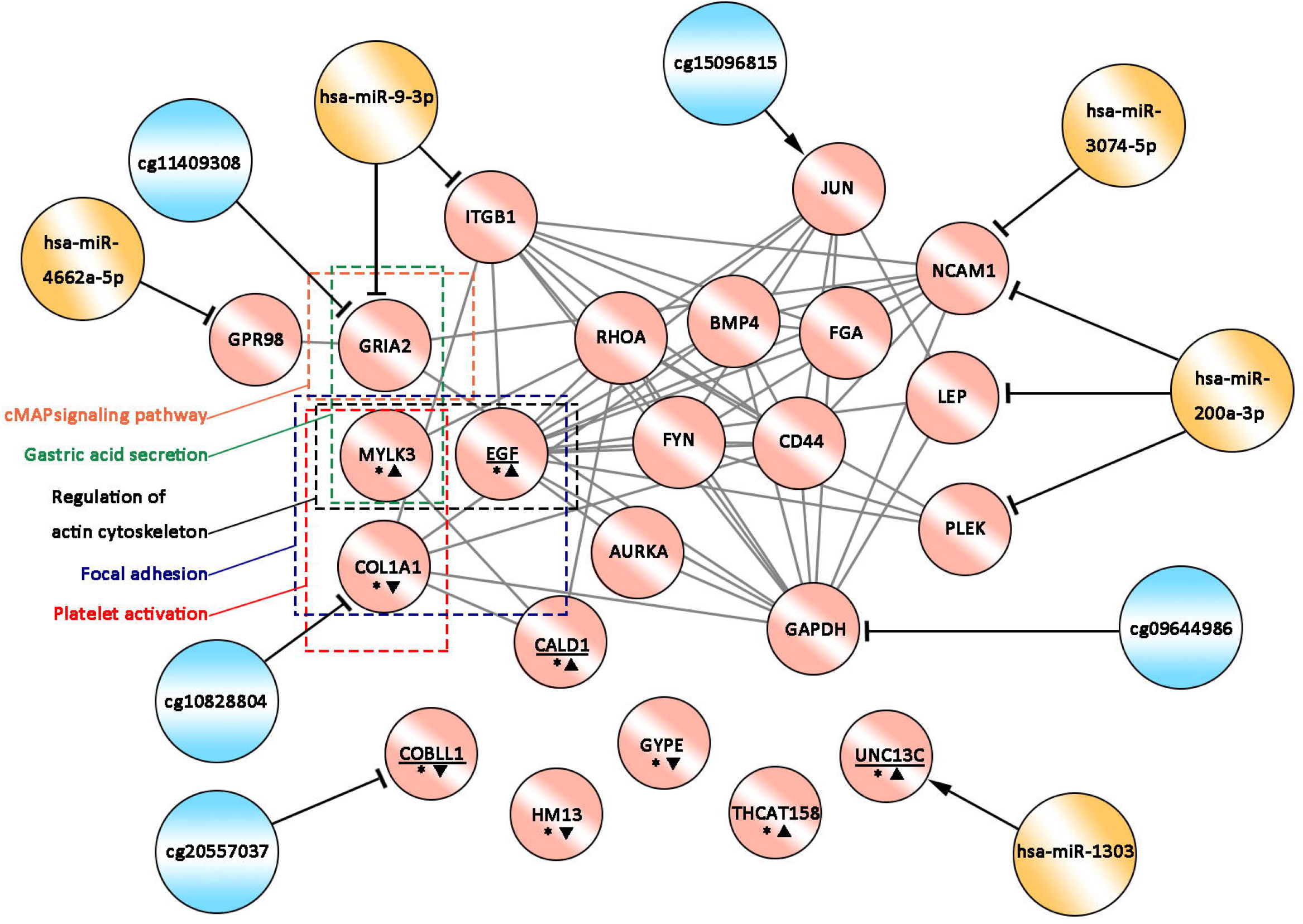
The predicted regulatory network of anti-HIV genes. Red circles denote the 25 anti-HIV genes based on prediction, blue circles denote DNA-methylation, and yellow circles denote DEMs. The lines with arrows pointing to red cycles denote promotion, while those with ‘T’ at the end denote inhibition. “*” in red circles denote the 12 anti-HIV genes with similar expression patterns both in EXP-Blood-HIV-Resistance and EXP-CD4-HIV-Resistance. Positive triangles next to “*” denote up-regulation, and inverted triangles denote down-regulation. The underlining of gene names denote the gene may be affected by the HIV Tat protein.

Previous analyses of EXP-Blood-HIV-Resistance has suggested that HIV-resistant people are always in a state of reduced immune activation [25]. It is reported that most genes are down-regulated in HIV-resistance pathways [25]. In contrast to previous results, our analyses of Blood-HIV-resistance showed that there are comparably more HIV-R up-regulated genes in most pathways. In the 24 identified HIV-resistant related pathways, only 7 pathways are with more HIV-R up-regulated than down-regulated genes, while the left 14 pathways are with more HIV-R down-regulated than up-regulated genes. There are only 2 pathways in common, Focal adhesion and Regulation of actin cytoskeleton, between the 24 pathways identified by us and the 15 pathways identified in the previous work [25]. In focal adhesion, previous work [25] found most genes are down-regulated, but our analysis showed that most genes (9/15) are up-regulated. In comparing the top 50 DEGs reported in the previous work and the identified 452 DEGs in our analysis, there are 11 genes in common and only 2 genes show consistent expression patterns. DPP4, referred as the most significant DEGs in [25], show no expression difference in our analysis. The previous work on CD4-HIV-Resistant proposed that the HIV-resistant CD4+ T cells are mostly silenced in gene expression [26]. This work reported 132 HIV-R DEGs, within which 124 were found to be in a low-expression in HIV-R cells. Based on these, it is proposed two related pathways, proteasome and the T cell receptor (TCR) signaling pathways, in which genes are identified mostly down-regulated [26]. Different from the previous results, our analysis found there were 4 down-regulated DEGs and 3 up-regulated DEGs in the previously reported HIV-resistant pathway, the T cell receptor (TCR) signaling (Figure S35). Moreover, the previous work [26] specially pointed out that IL-5, a member of human Interleukin, is down-regulated in HIV-resistant CD4+ T cells. In contrast, our analysis showed that this gene is up-regulated. The difference between our results and the previous work [26] may be due to the exclusion of low quality samples in the sample quality control of our analysis. CCR5 is a validated anti-HIV gene [53], whose mutations have a higher frequency in the Caucasian population [54], This gene is not included in our results, so did previous records [25, 26].

Two identified HIV-resistant genes have been previously reported. They are NCAM1 and RhoA. NCAM1, neural cell adhesion molecule, encodes the cell adhesion protein, a member of human immunoglobulins [55], and mainly expresses in natural killer cells (NK) [56], T cells, B cells and dendritic cells [57, 58]. These cells play important roles in immune surveillance. NCAM1 can also be detected in neural and mesenchymal stem cells [59]. It functions in the development of the nervous system by regulating neurogenesis, neurite outgrowth, and cell migration [60–62], It has been reported that NK cells of HIV exposure show a special activated state, e.g. the promoted expression of CD107a, IFN-γ, CD57 and CD56 [63]. These indicated that NCAM1 play an important role in the resistance to HIV infection. RhoA, Ras Homolog Family Member A, encodes a GTPase, a member of Rho family. RhoA functions importantly in multiple cell activities, e.g., cell growth, cell transformation and cytoskeleton regulation [64–66]. It has been reported that RhoA is capable of regulating and inhibiting the replication of HIV-1 through novel effector activity [67],

For the 23 newly identified HIV-resistance related genes, most (14/23) are in relationship with the lifecycle of tumor cells, such as AURKA [68–70], BMP4[24, 71, 72] and CALD1 [25, 66, 73] (Table S40). Moreover, five HIV-resistance genes are neurological function related (Table S8). It can be inferred that investigation on cancer and neurological diseases may provide us novel hints for the therapy and the prevention of HIV/AIDS. In past decades, the limited successfully cured AIDS cases all have acute myeloid leukemia (AML) and the patients were all treated by hemopoietic stem cell transplantation (HSCT) [74, 75] [76] [77], As the blood cancer was treated, the HIV in the patient’s body was also eliminated. However, the detailed mechanism is still a mystery.

## Author Contributions

G.L. collected, analyzed and compiled the data. G.L., C.Z., L.S. and Z.Z. took part in the analysis. Z.Z. conceived the idea. Z.Z. and L.S. coordinated the project. Z.Z., G.L. and L.S. wrote the manuscript.

## Acknowledgments

We gratefully acknowledge the submitting and the originating laboratories where genetic sequence data were generated and shared via NCBI. This work was supported by the Fundamental Research Funds for the Central Universities (2019CDYGYB024) to LS and the National Natural Science Foundation of China (31200941) to ZZ.

