## Supplementary Figures for "Identification of regulatory networks associated with anti-HIV/AIDS genes via integrated transcriptome, epigenome and proteome analyses"

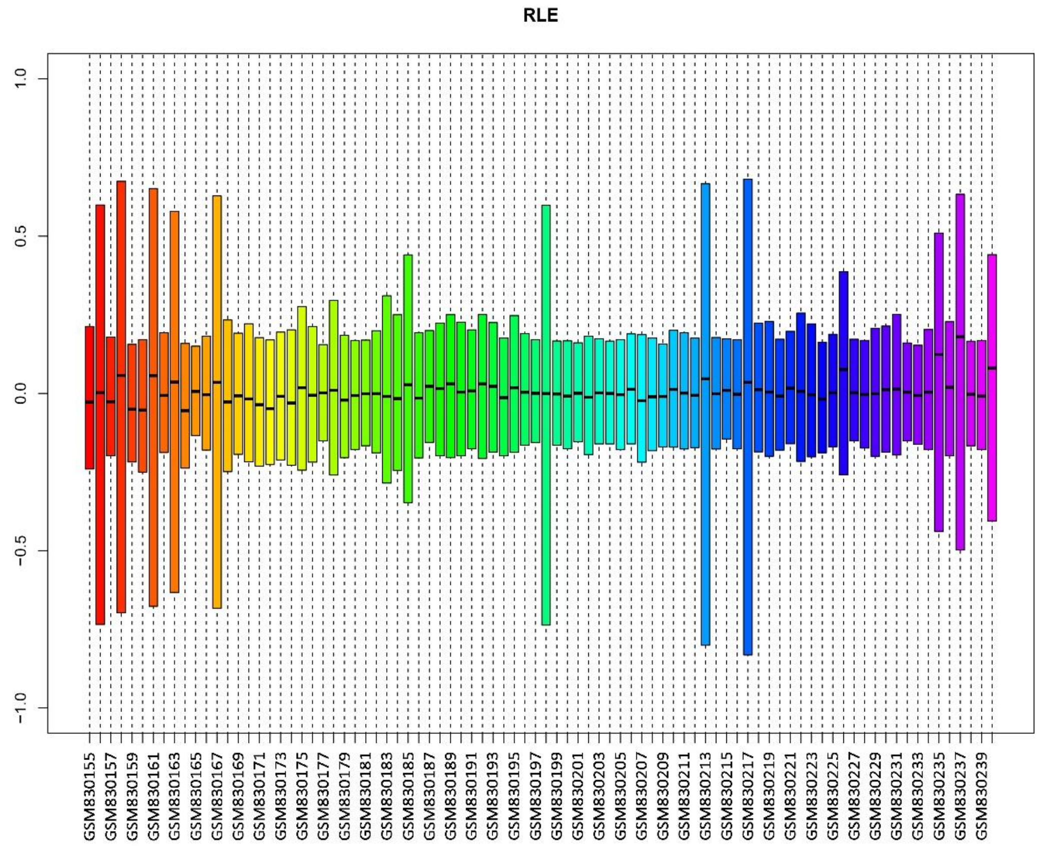


Figure S1. The RLE box plot of samples in the transcriptome data set EXP-Blood-HIV-Resistance. This figure is produced by the affy package for quality control.

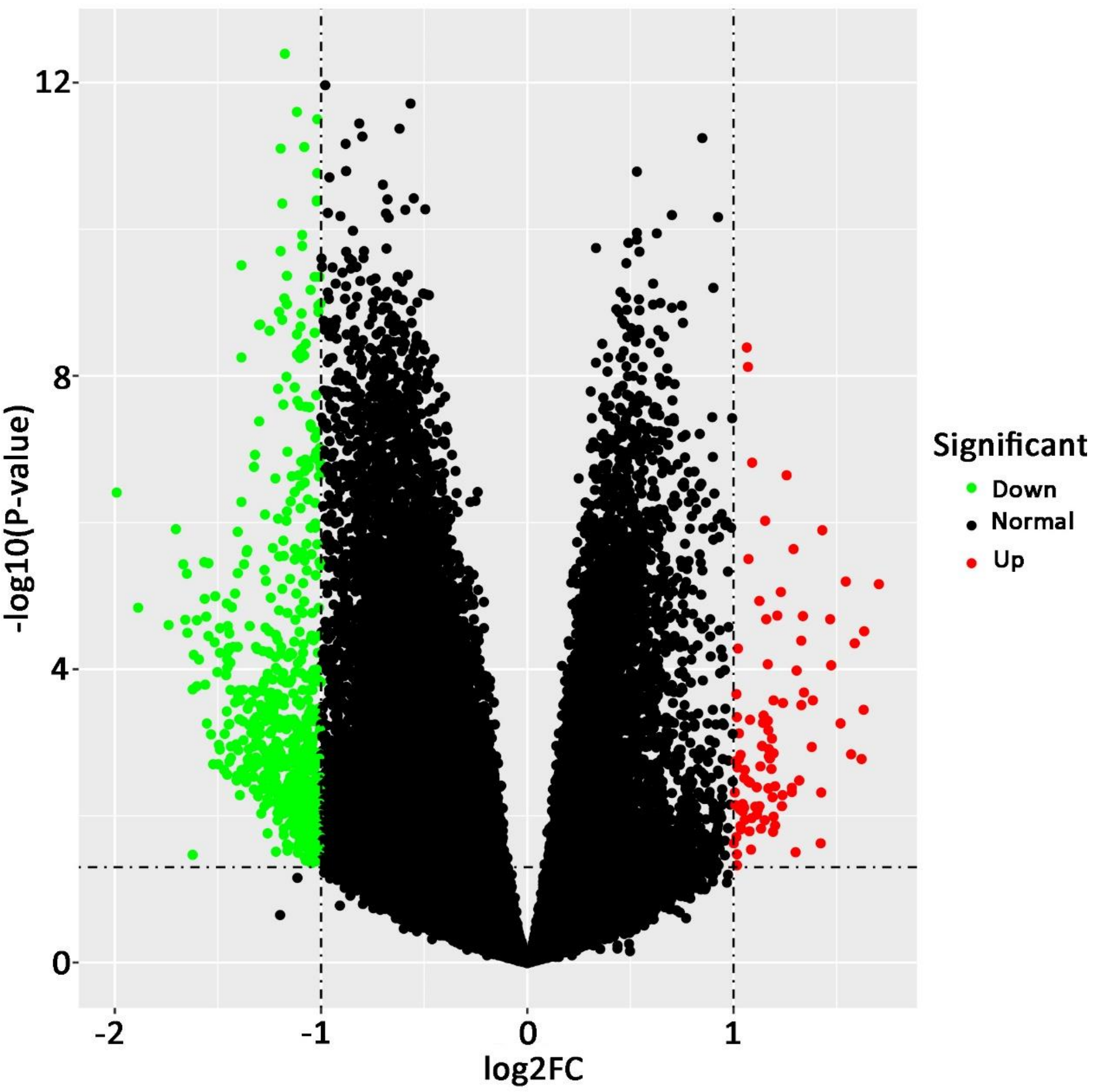


Figure S2. The volcano plot of gene expression changes between HIV-R and HIV-N-C in EXP-Blood-HIV-Resistant.

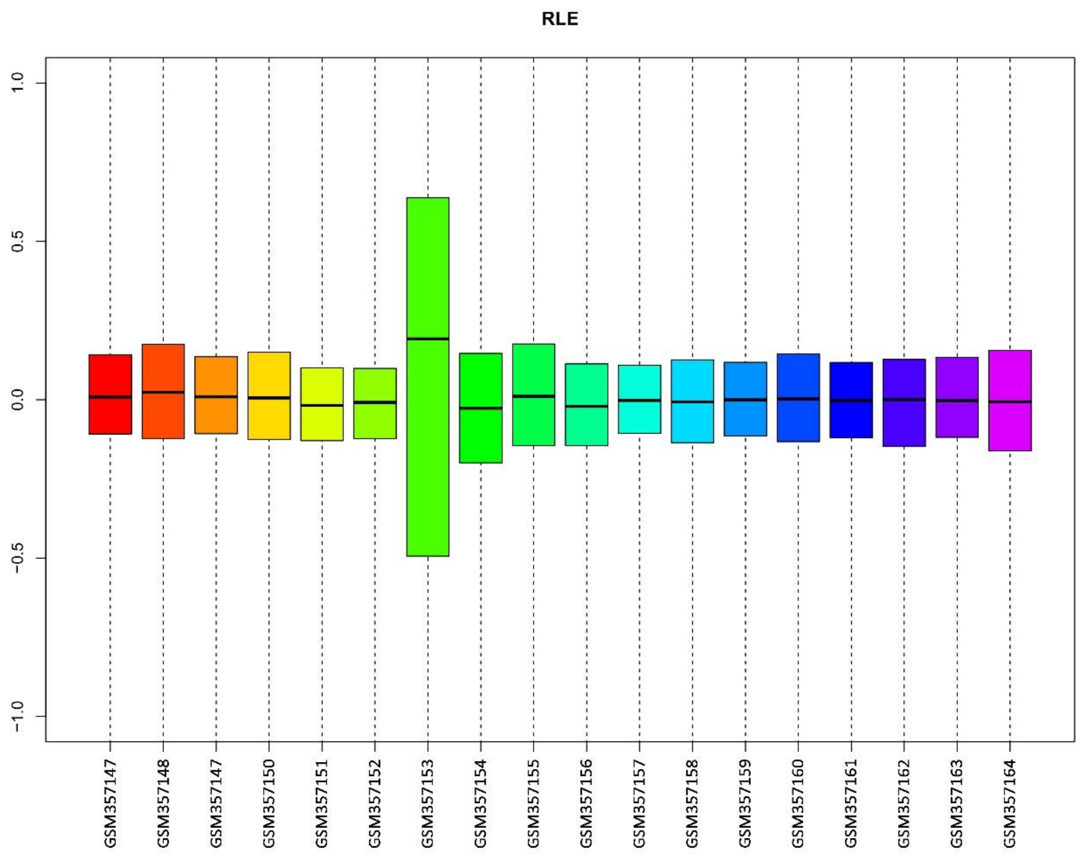


Figure S3. The RLE box plot of samples in the transcriptome data set EXP-CD4-HIV-Resistance. Other legends follow Figure S1.

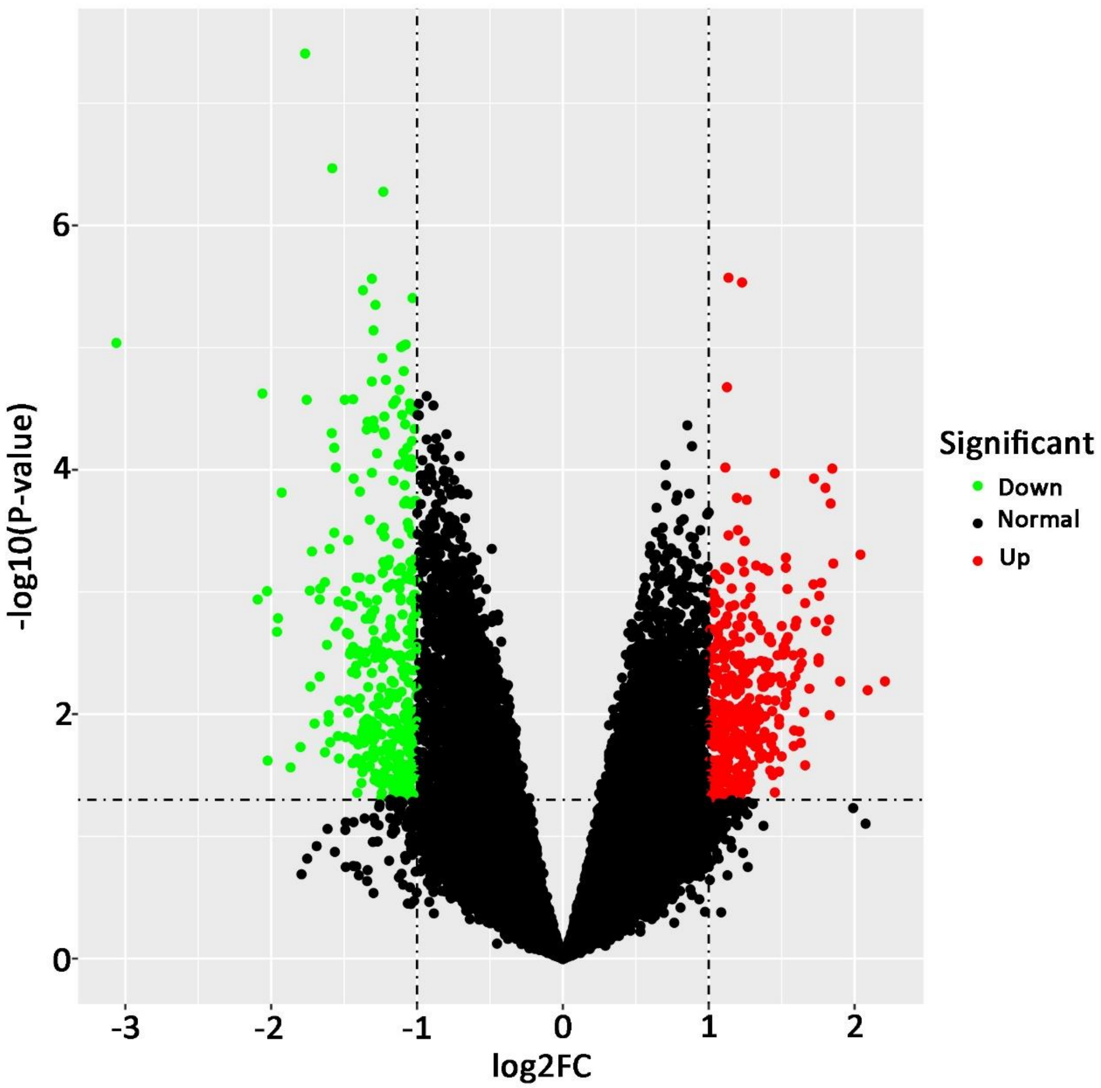


Figure S4. The volcano plot of gene expression changes between HIV-R and HIV-N-C in EXP-CD4-HIV-Resistant.

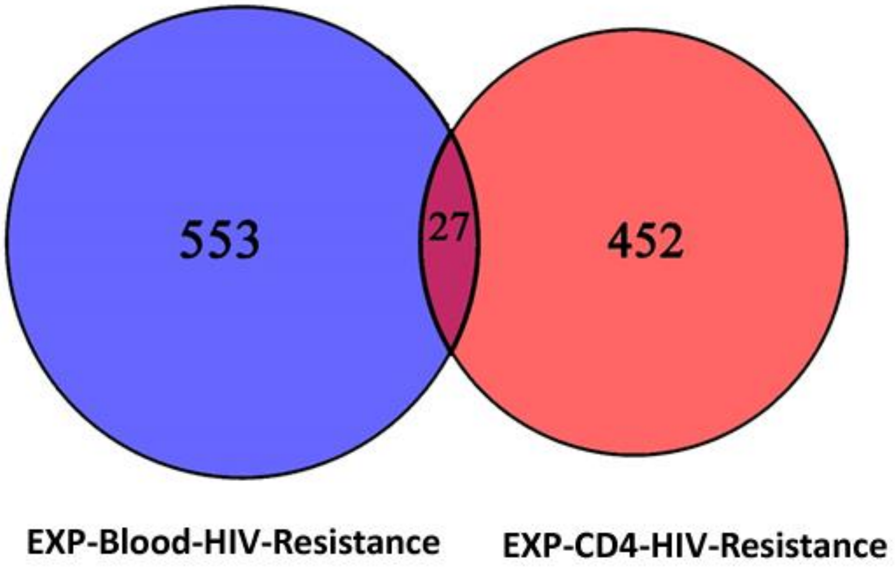


Figure S5. A Venn plot showing the overlap of DEGs resulted from EXP-Blood-HIV-Resistance and EXP-CD4-HIV-Resistance.

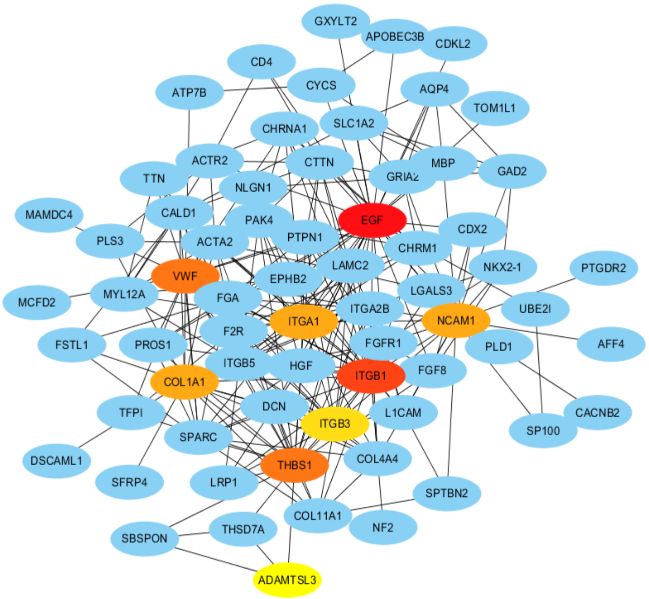


Figure S6. The PPI network of DEGs of EXP-Blood-HIV-Resistance using the Degree algorithm.

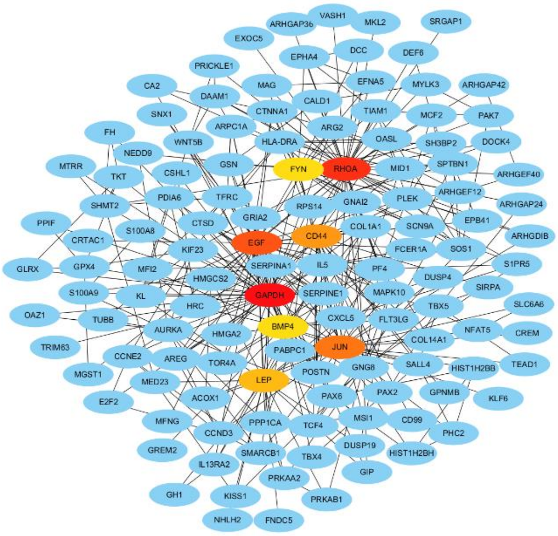


Figure S7. The PPI network of DEGs of EXP-CD4-HIV-Resistance using the Degree algorithm.

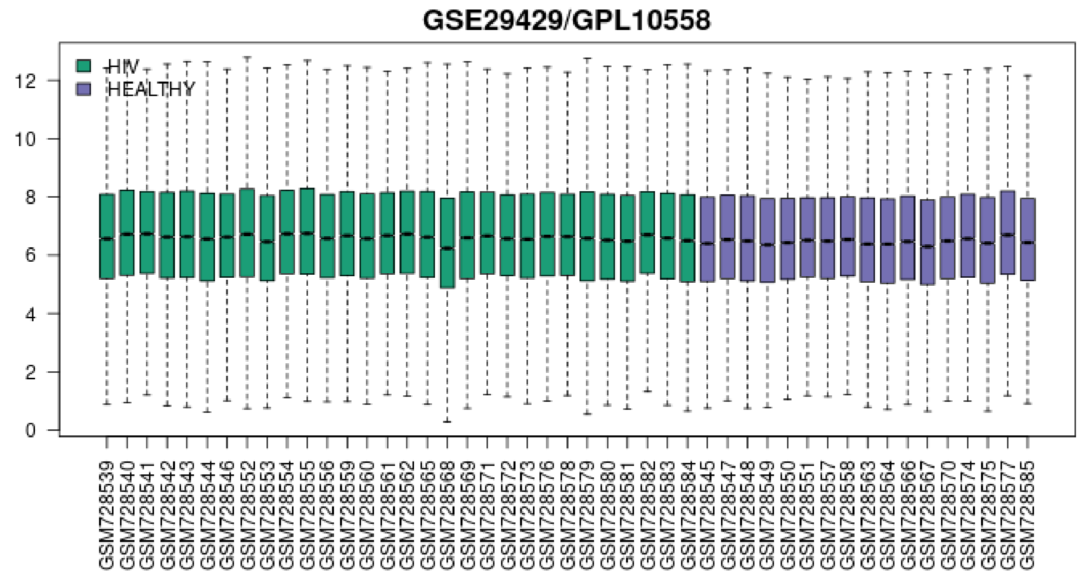


Figure S8. Box plot of the transcriptome dataset EXP-Blood-HIV-Infection. This figure is produced by GEO2R for quality control. X axis are samples. Y are median-centered values indicative if data are normalized and cross-comparable.

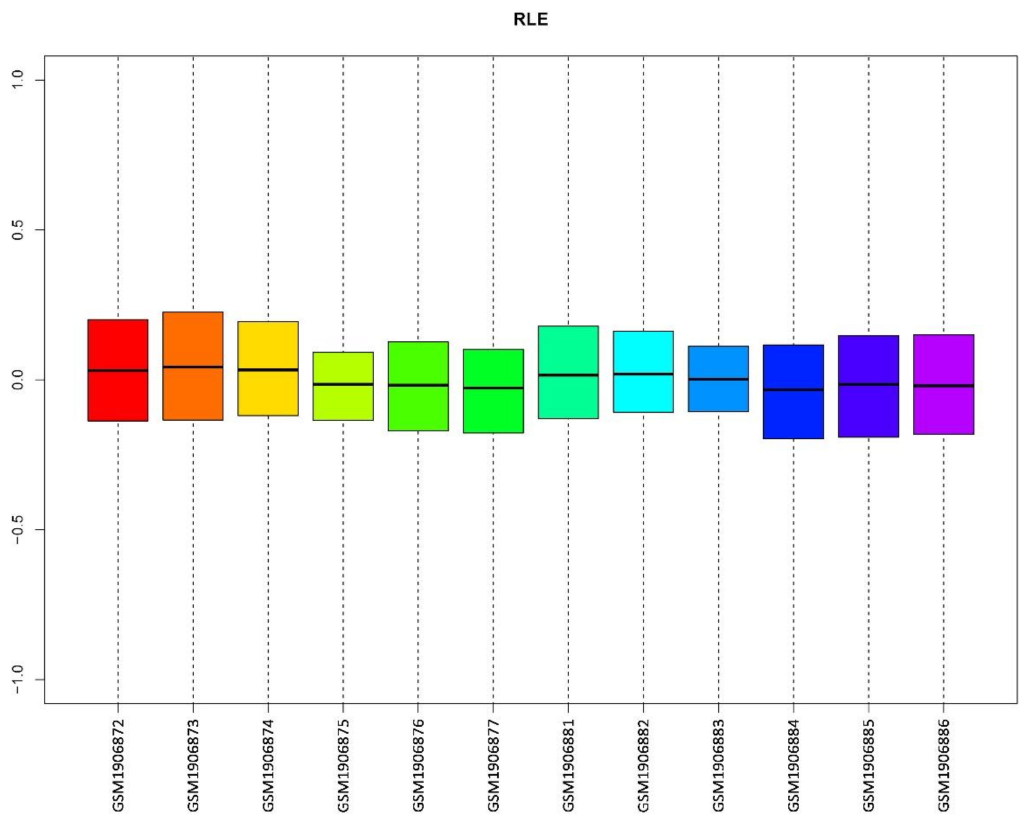


Figure S9. The RLE box plot of samples in the transcriptome data set EXP-CD4-HIV-Infection. Other legends follow Figure S1.

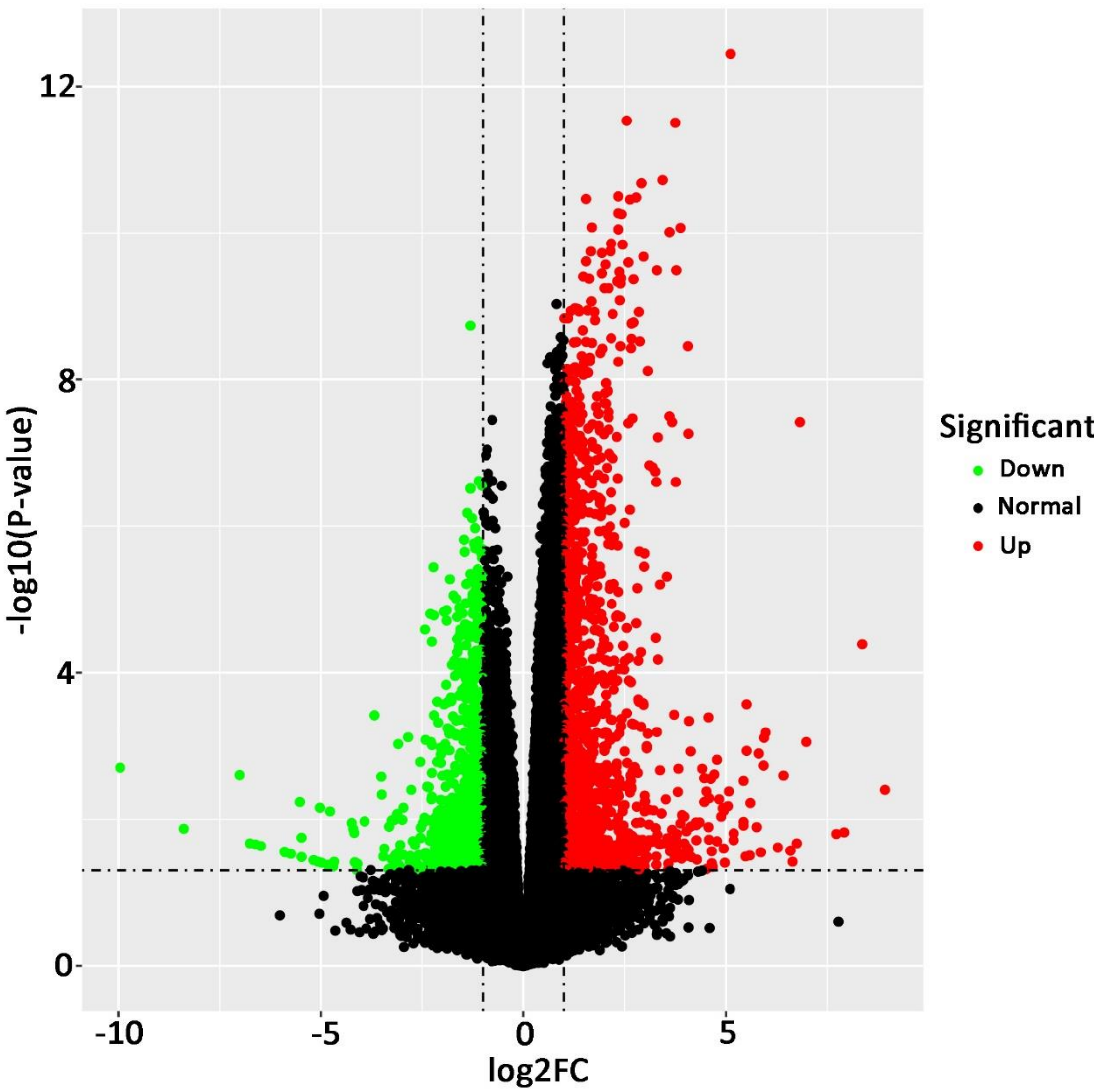


Figure S10. The volcano plot of gene expression changes between HIV+ and HIV- in EXP-Blood-HIV-Infection.

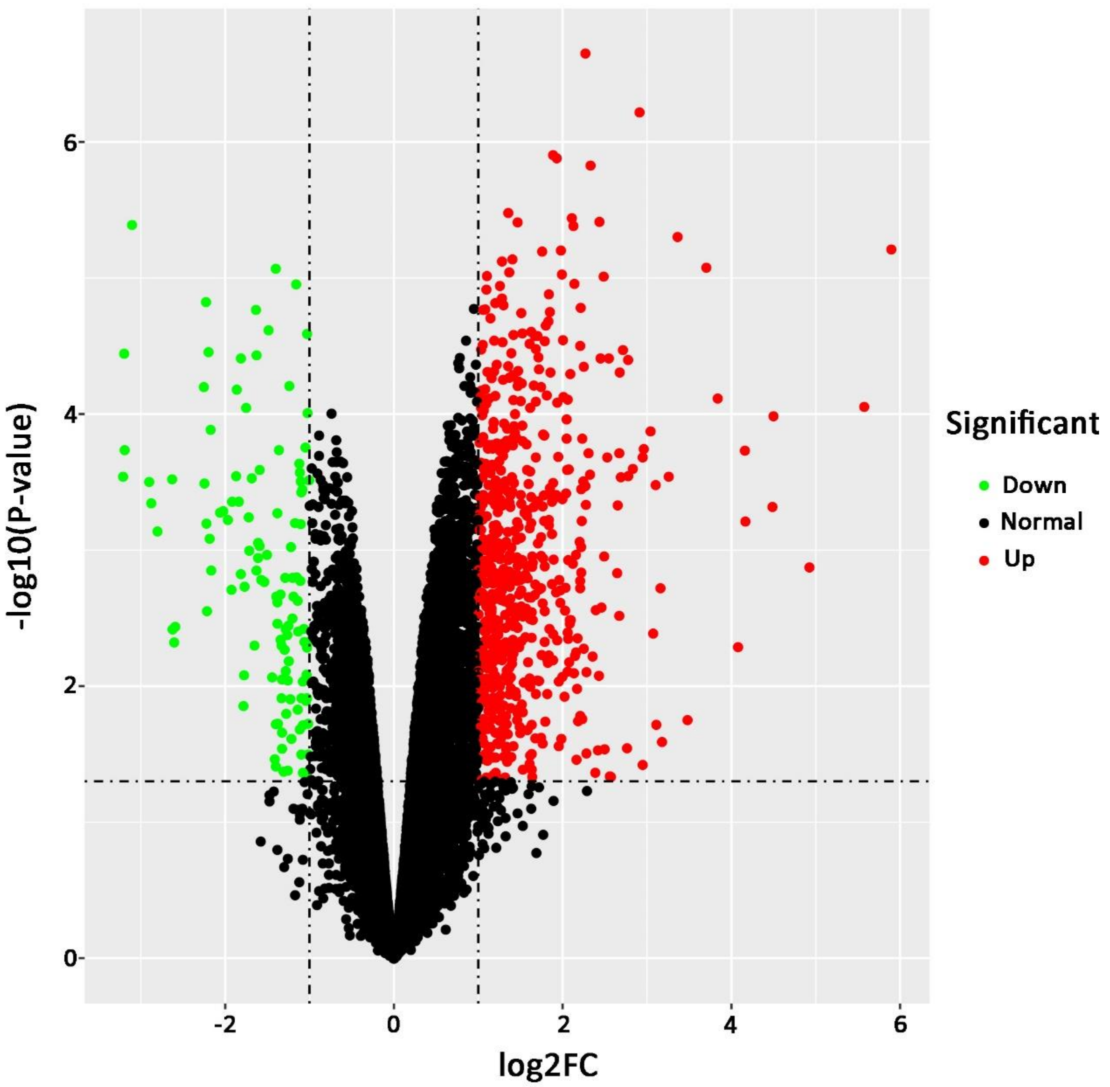


Figure S11. The volcano plot of gene expression changes between HIV+ and HIV- in EXP-Blood-HIV-Infection (Naive CD4+ T cell).

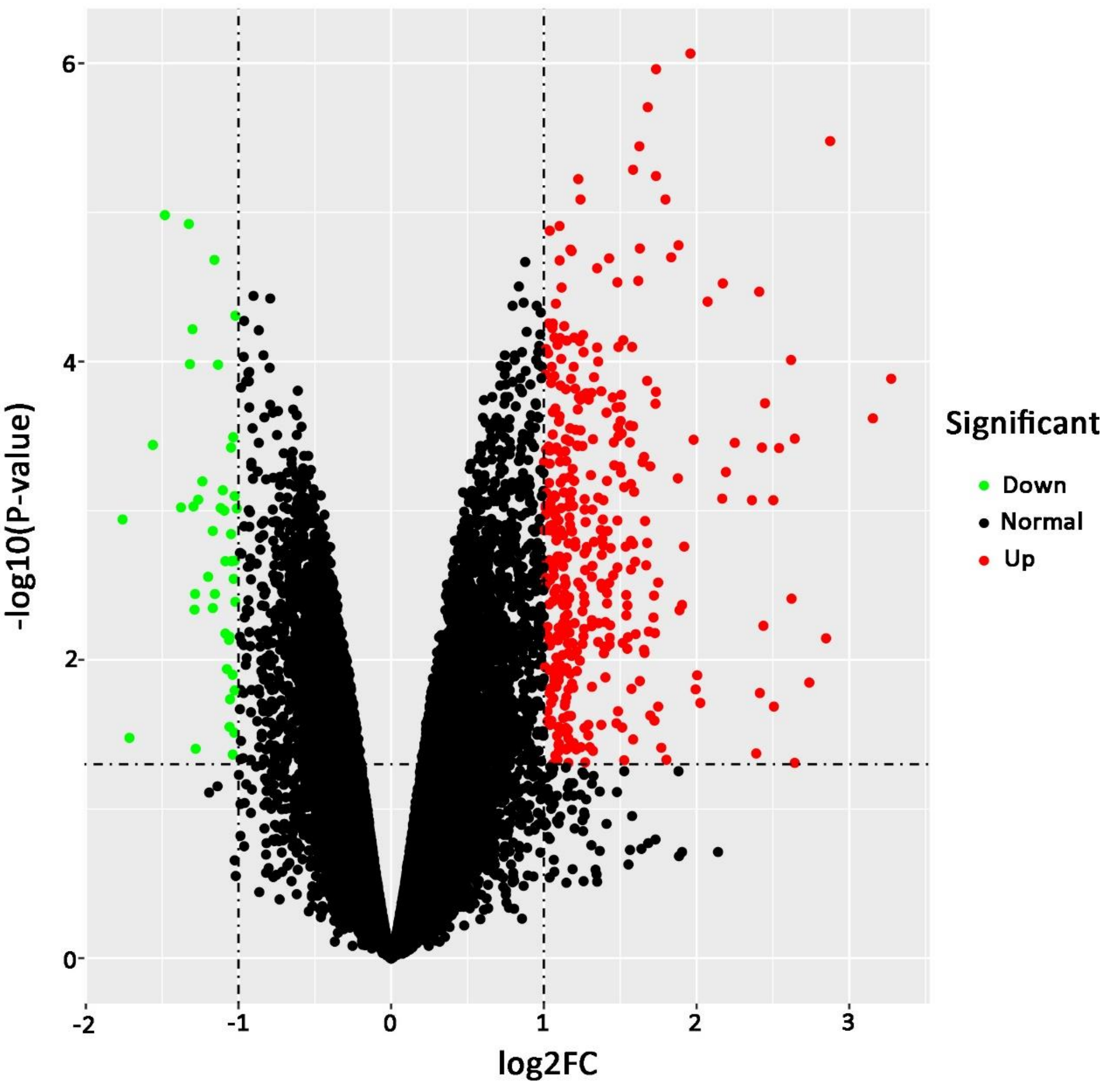


Figure S12. The volcano plot of gene expression changes between HIV+ and HIV- in EXP-Blood-HIV-Infection (Central memory CD4+ T cells).

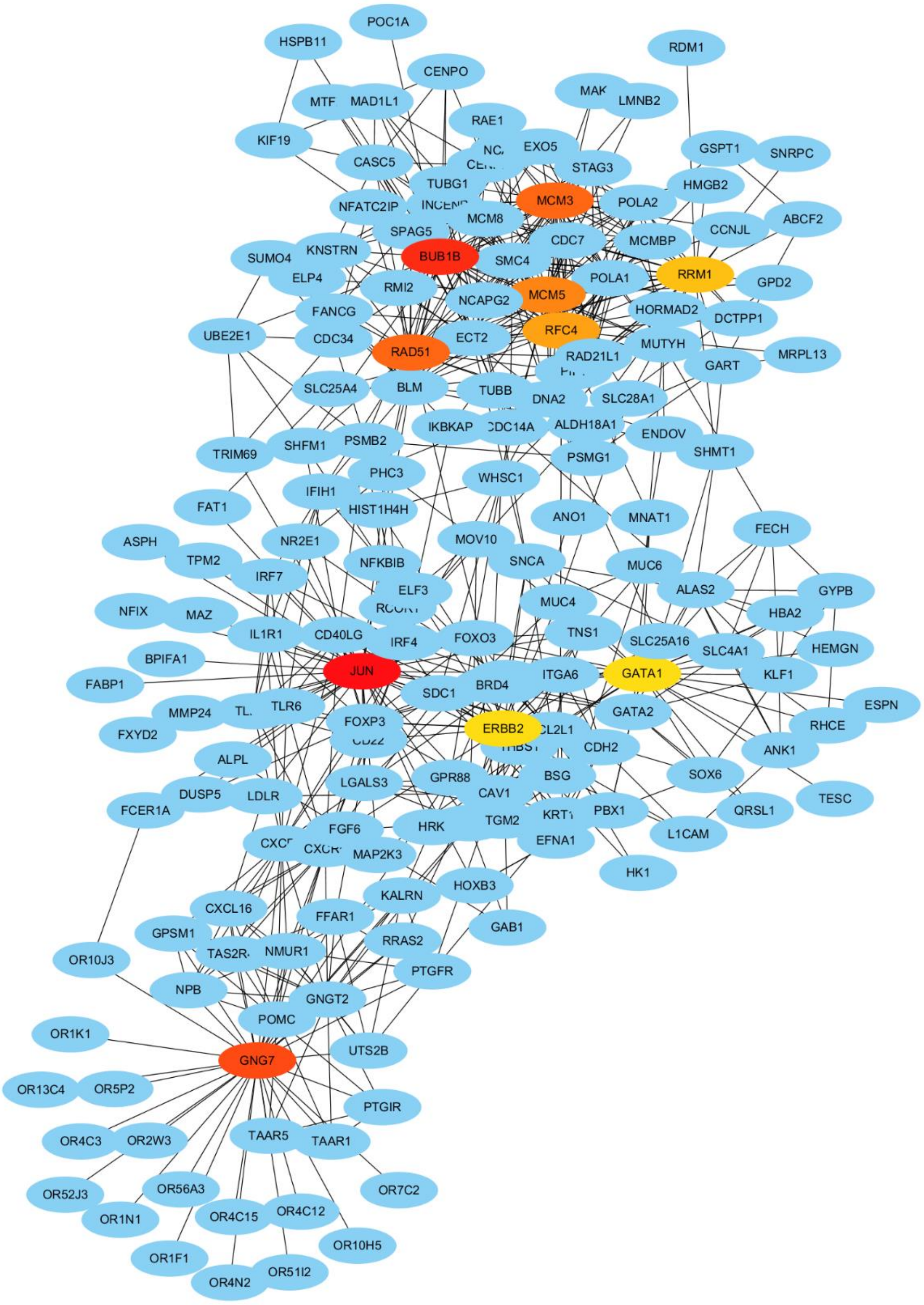


Figure S13. The PPI network of DEGs of EXP-Blood-HIV-Infection using the Degree algorithm.

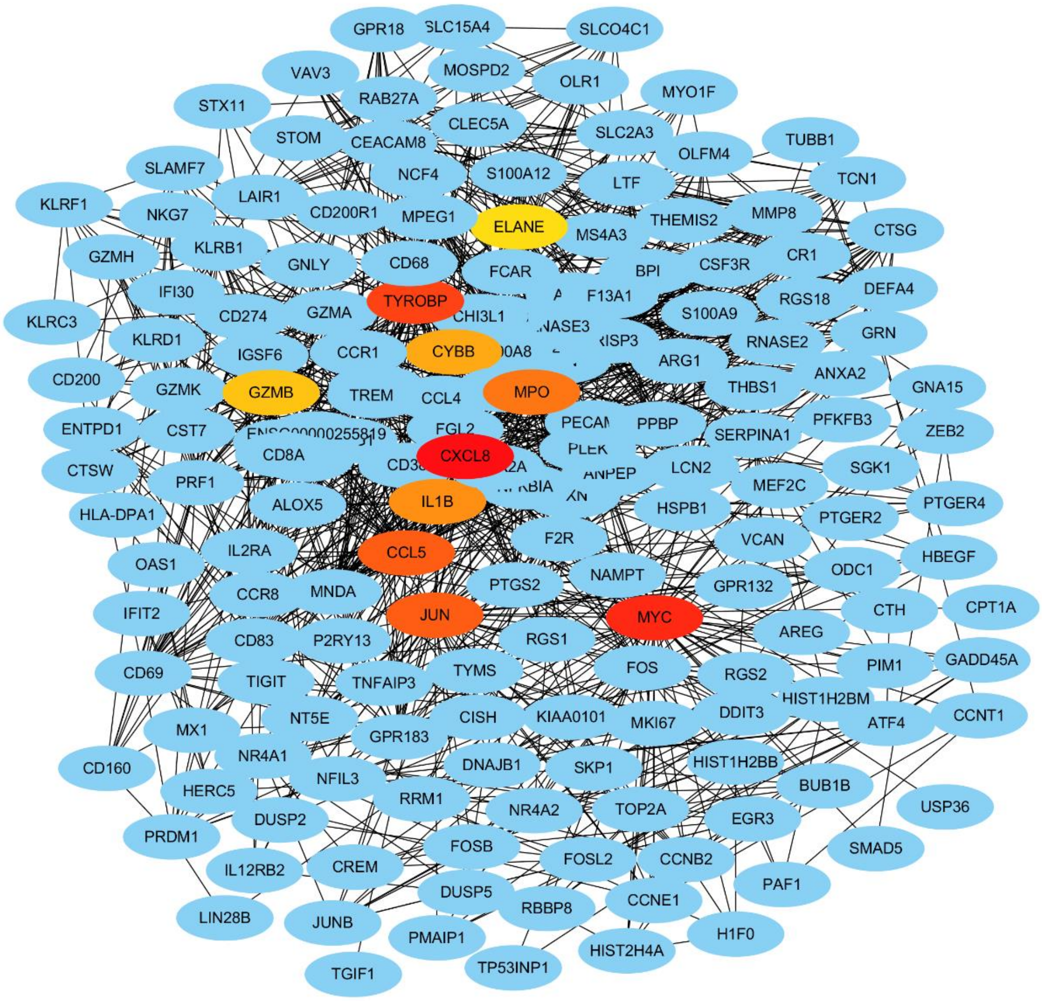


Figure S14. The PPI network of DEGs of EXP-CD4-HIV-Infection using the Degree algorithm.

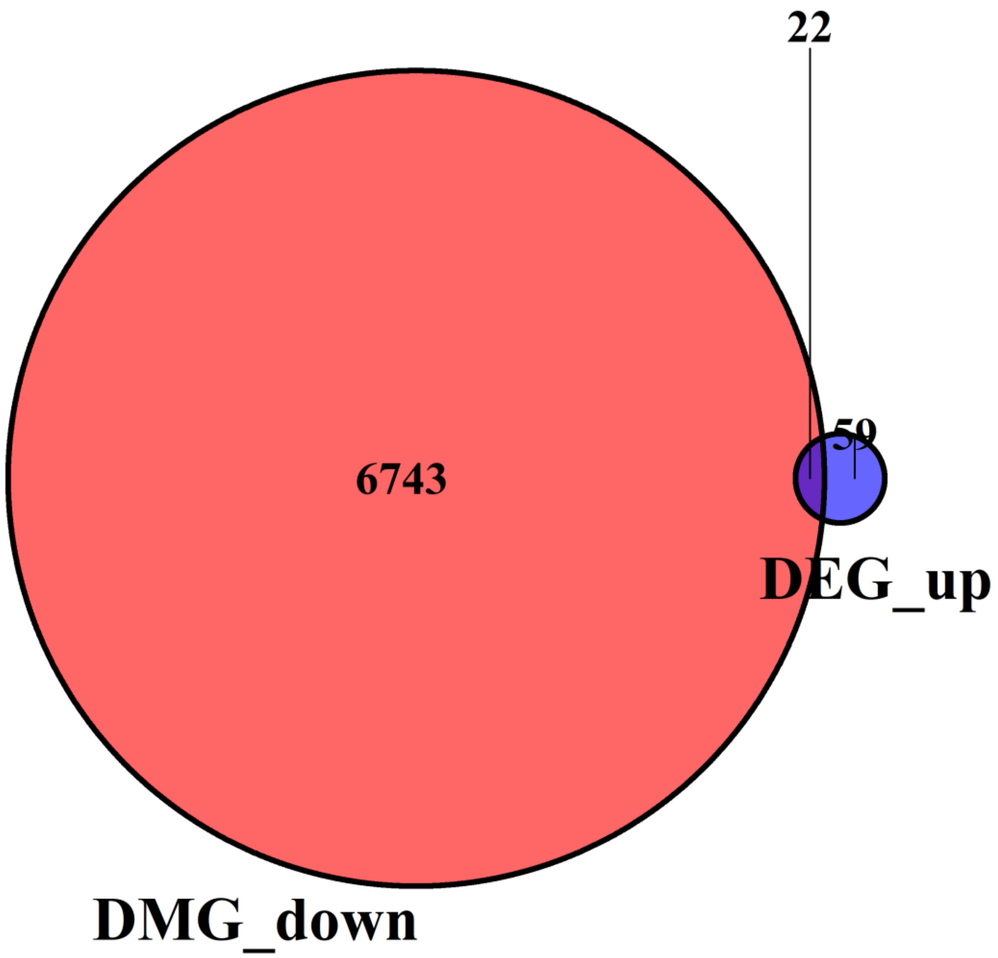


Figure S15. The Venn plot of hypomethylated up-regulated genes. The overlap of the HIV-R up-regulated DEGs and the hypomethylated DMGs.

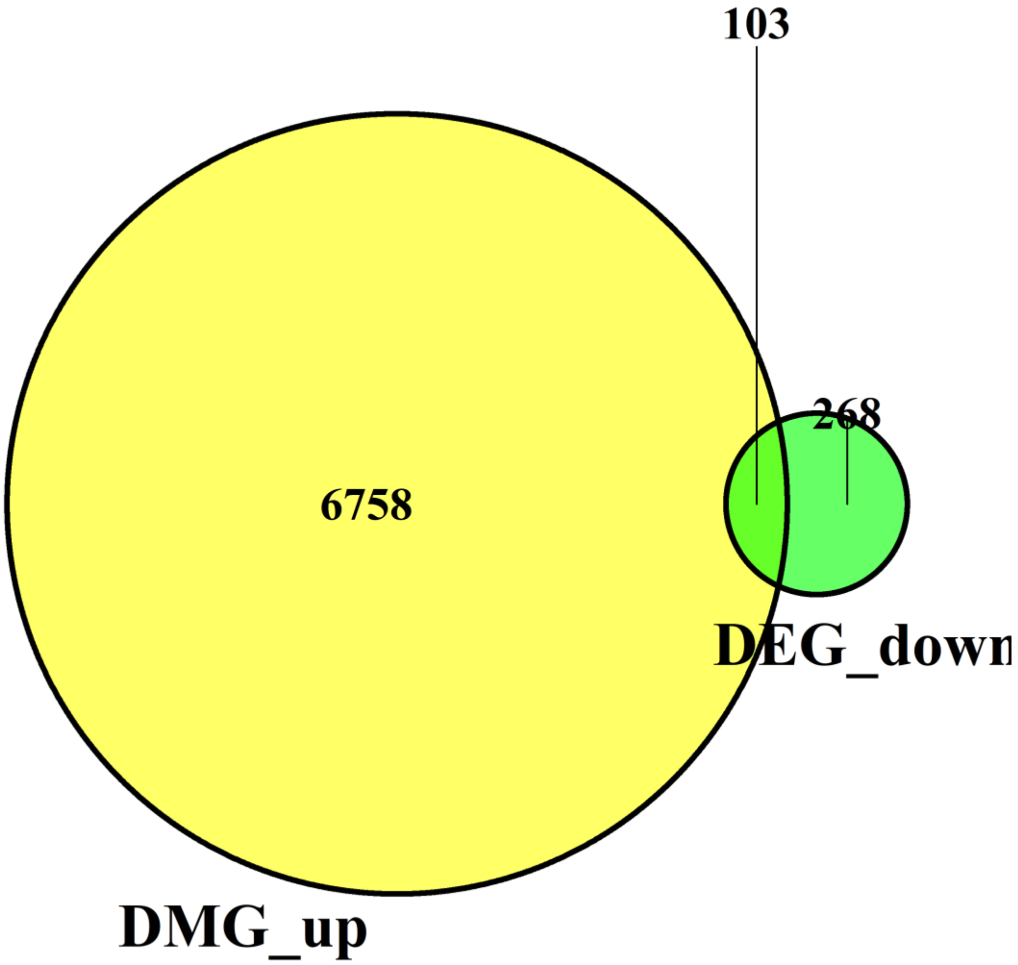


Figure S16. The Venn plot of hypermethylated down-regulated genes. The overlap of the HIV-R down-regulated DEGs and the hypermethylated DMGs.

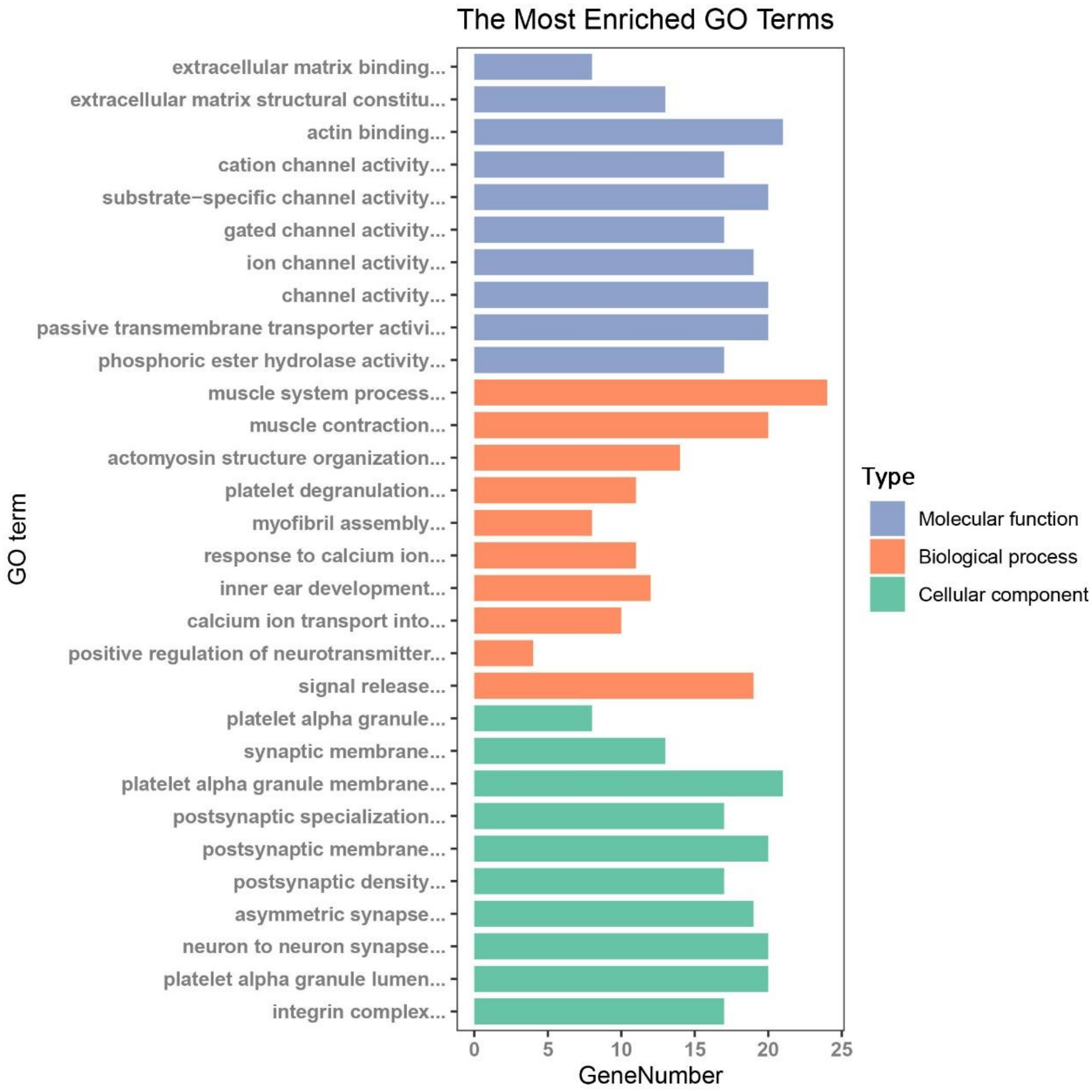


Figure S17. The GO enrichment analysis plot of DEGs in EXP-Blood-HIV-Resistance.

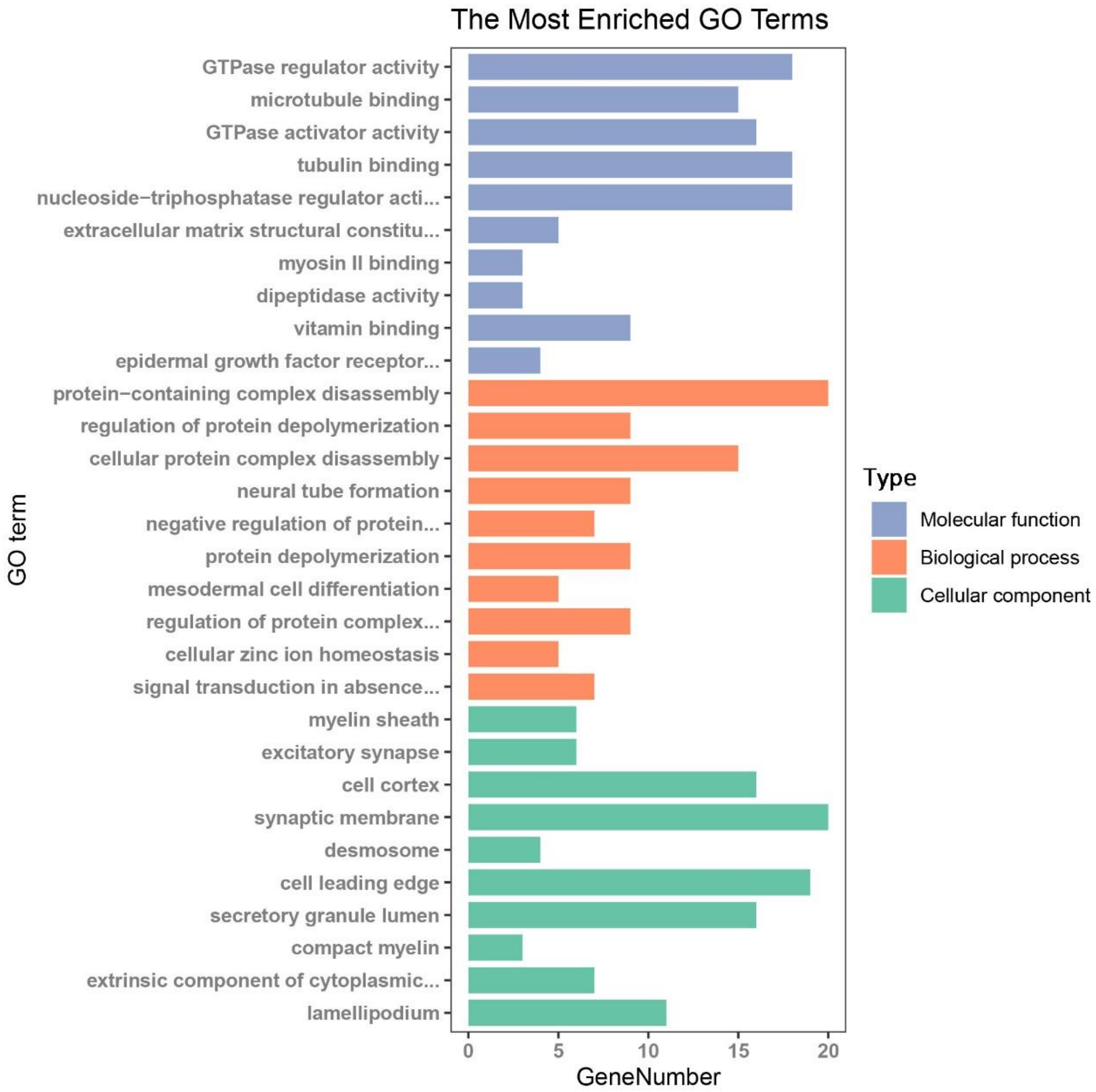


Figure S18. The GO enrichment analysis plot of DEGs in EXP-CD4-HIV-Resistance.

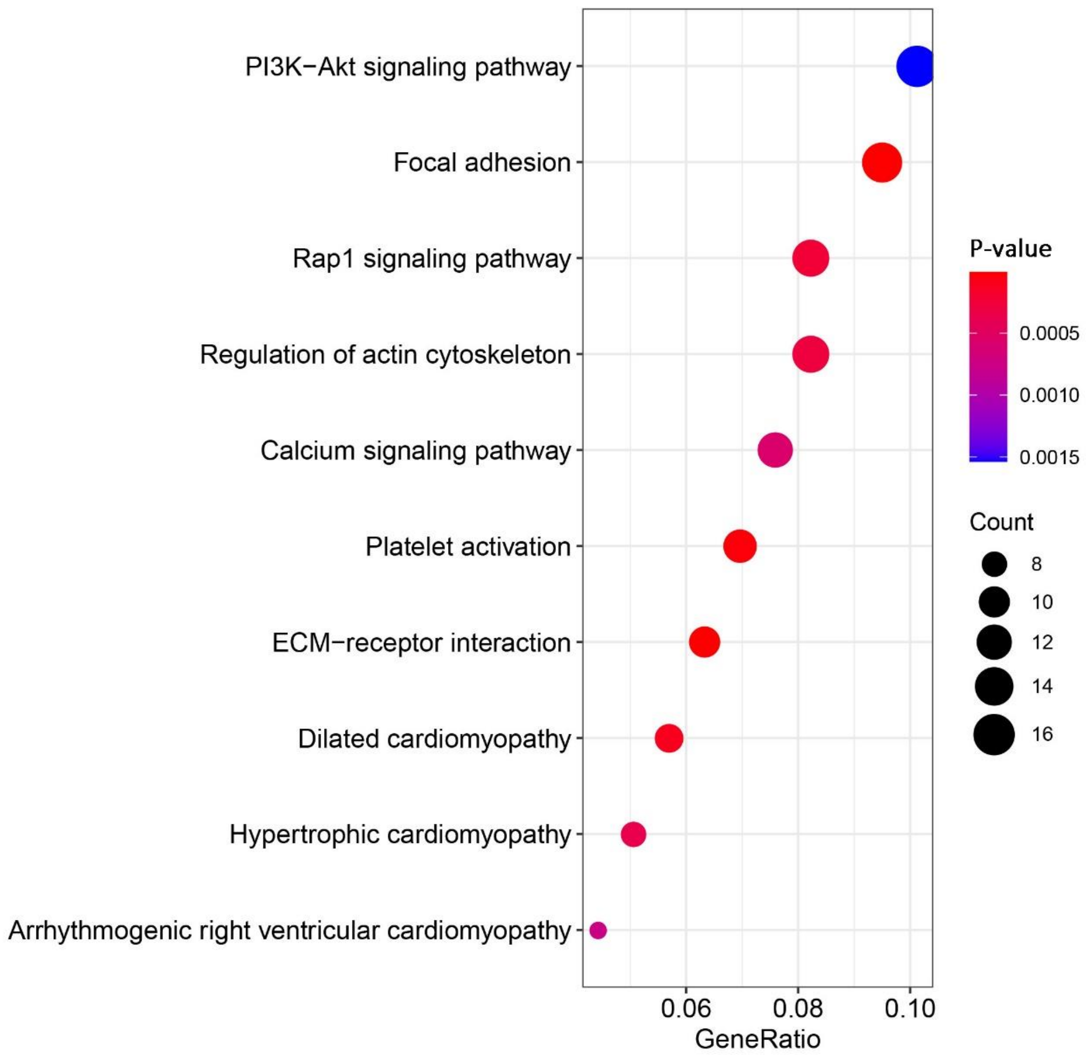


Figure S19. The pathway enrichment analysis plot of the DEGs in EXP-Blood-HIV-Resistance.

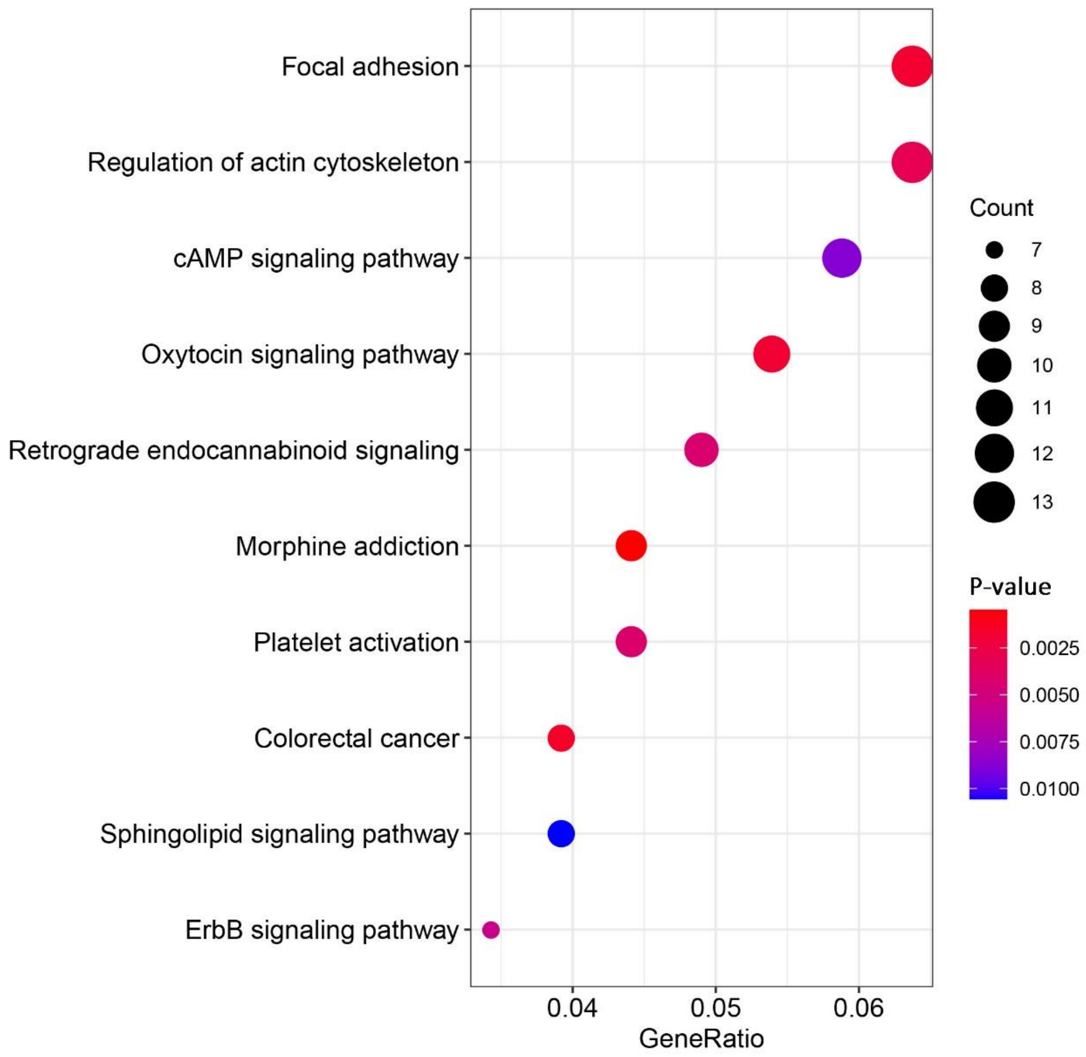


Figure S20. The pathway enrichment analysis plot of the DEGs in EXP-CD4-HIV-Resistance.

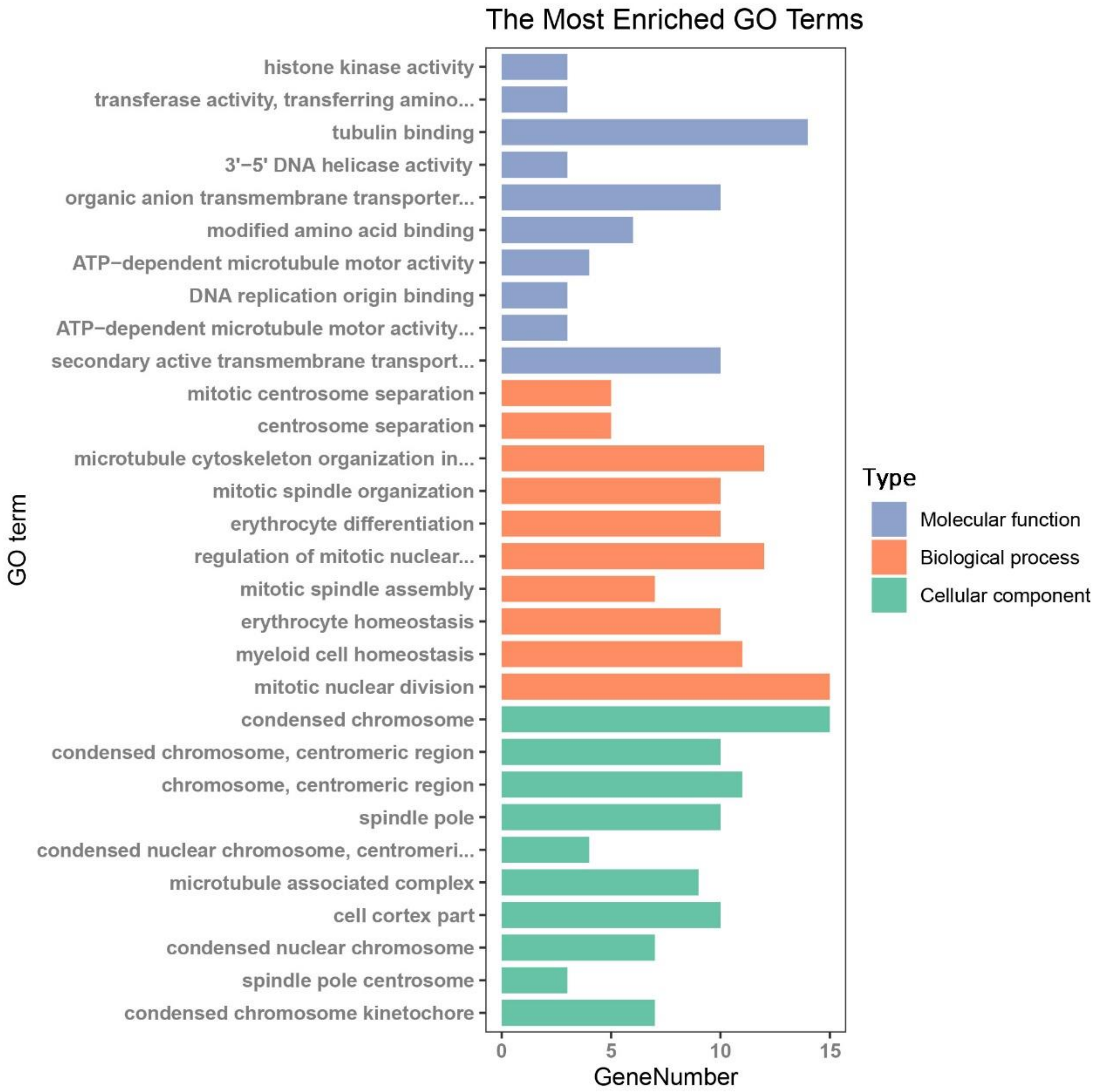


Figure S21. The GO enrichment analysis plot of the DEGs in EXP-Blood-HIV-Infection.

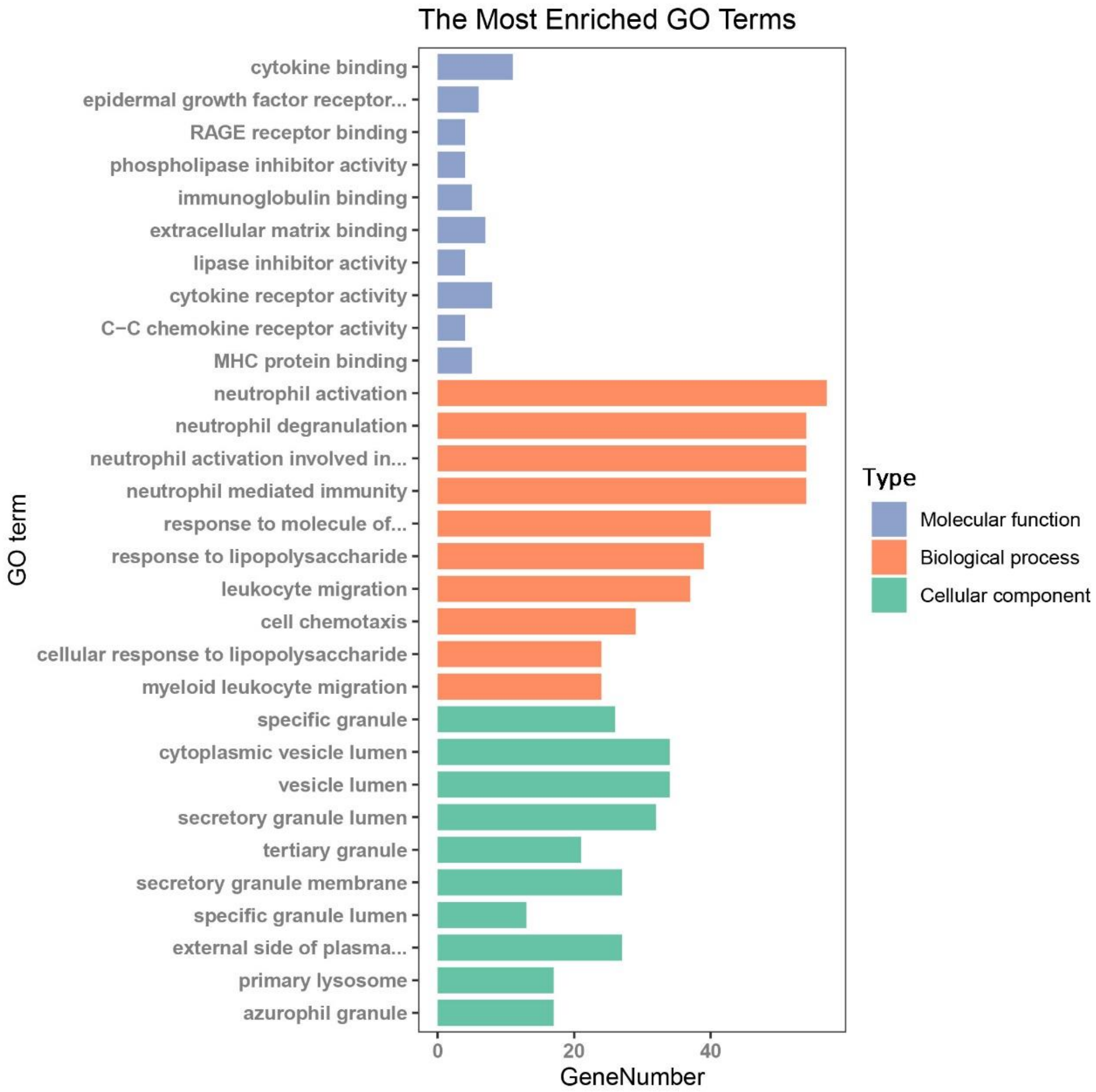


Figure S22. The GO enrichment analysis plot of the DEGs in EXP-CD4-HIV-Infection.

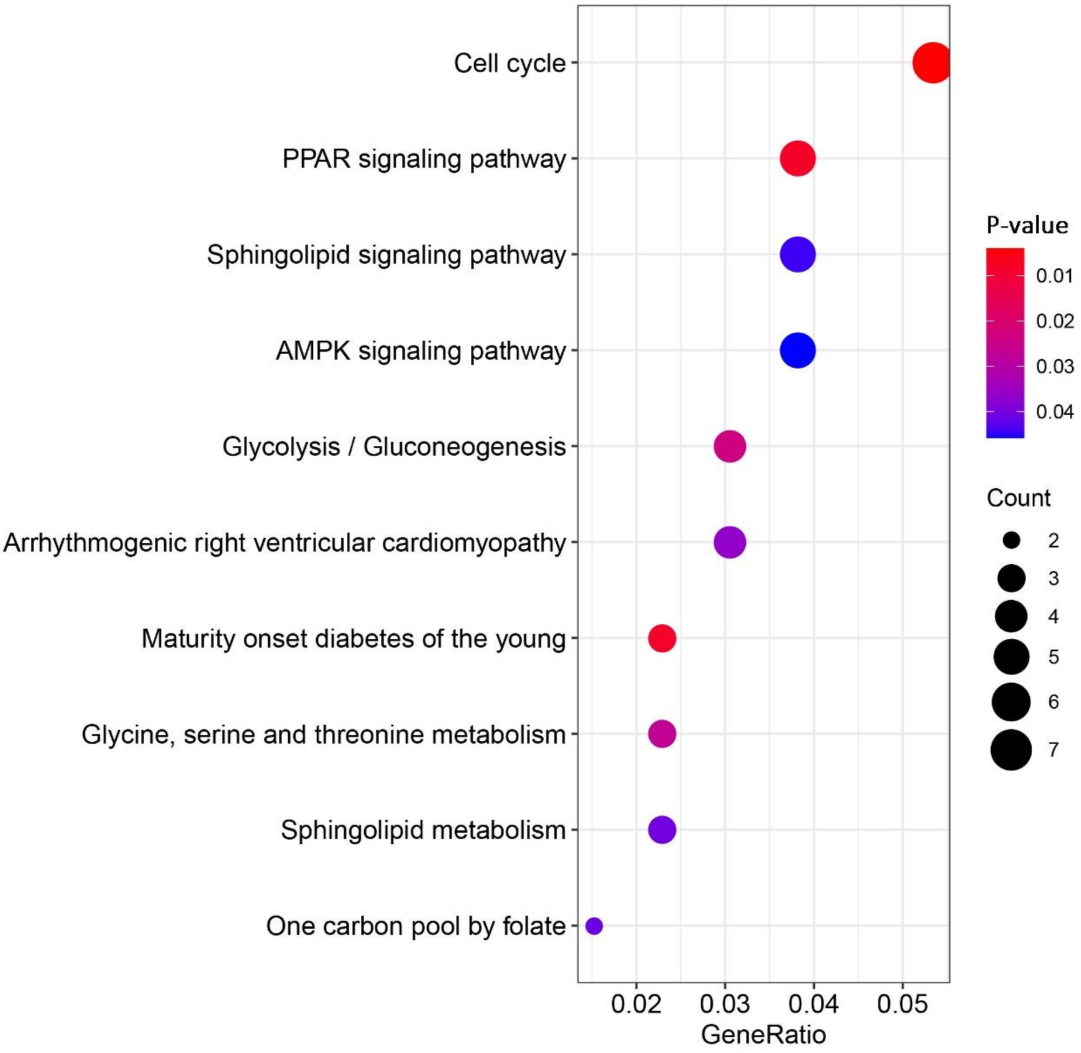


Figure S23. The pathway enrichment analysis plot of the DEGs in EXP-Blood-HIV-Infection.

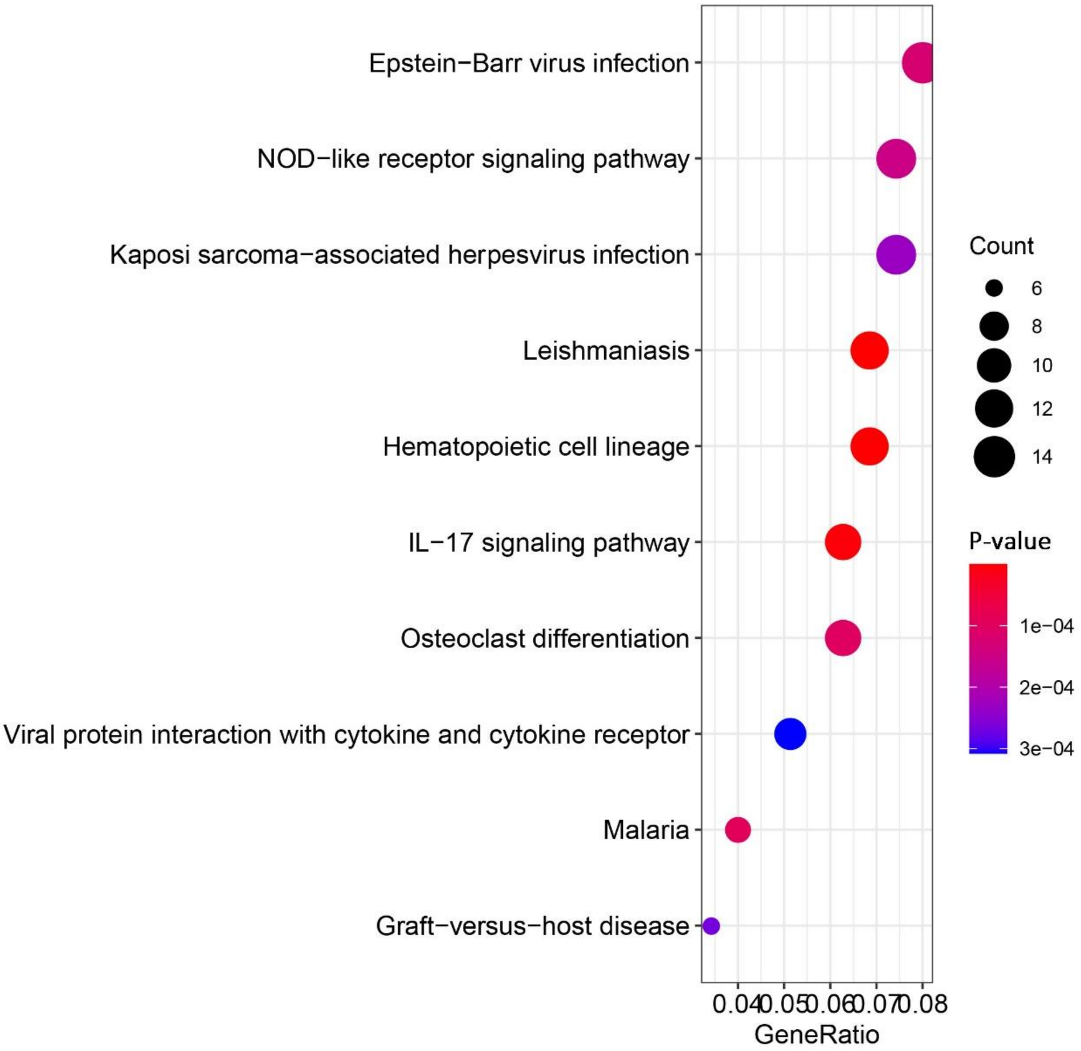


Figure S24. The pathway enrichment analysis plot of the DEGs in EXP-CD4-HIV-Infection.

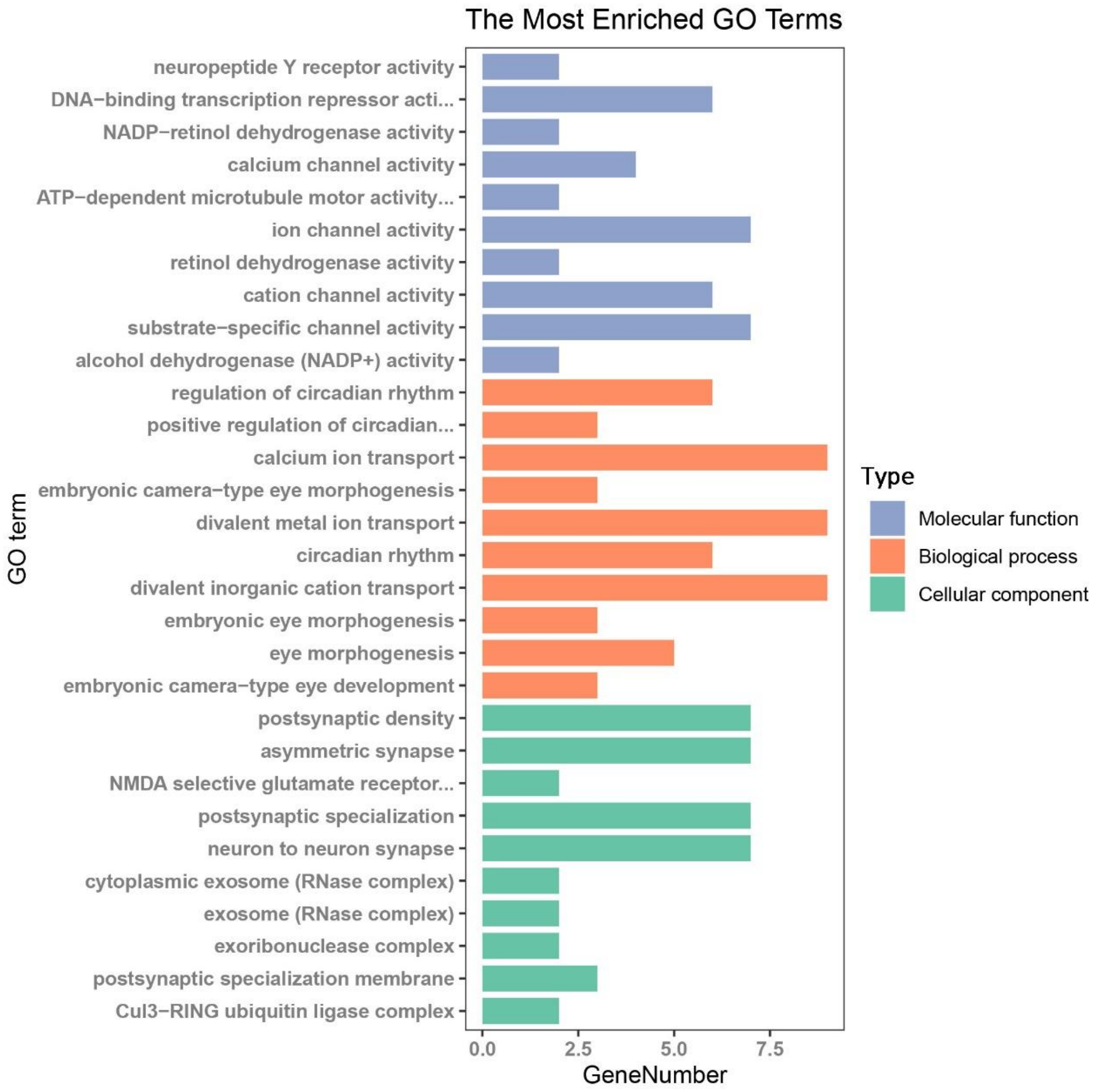


Figure S25. The GO enrichment analysis plot of the hypermethylated down-regulated genes.

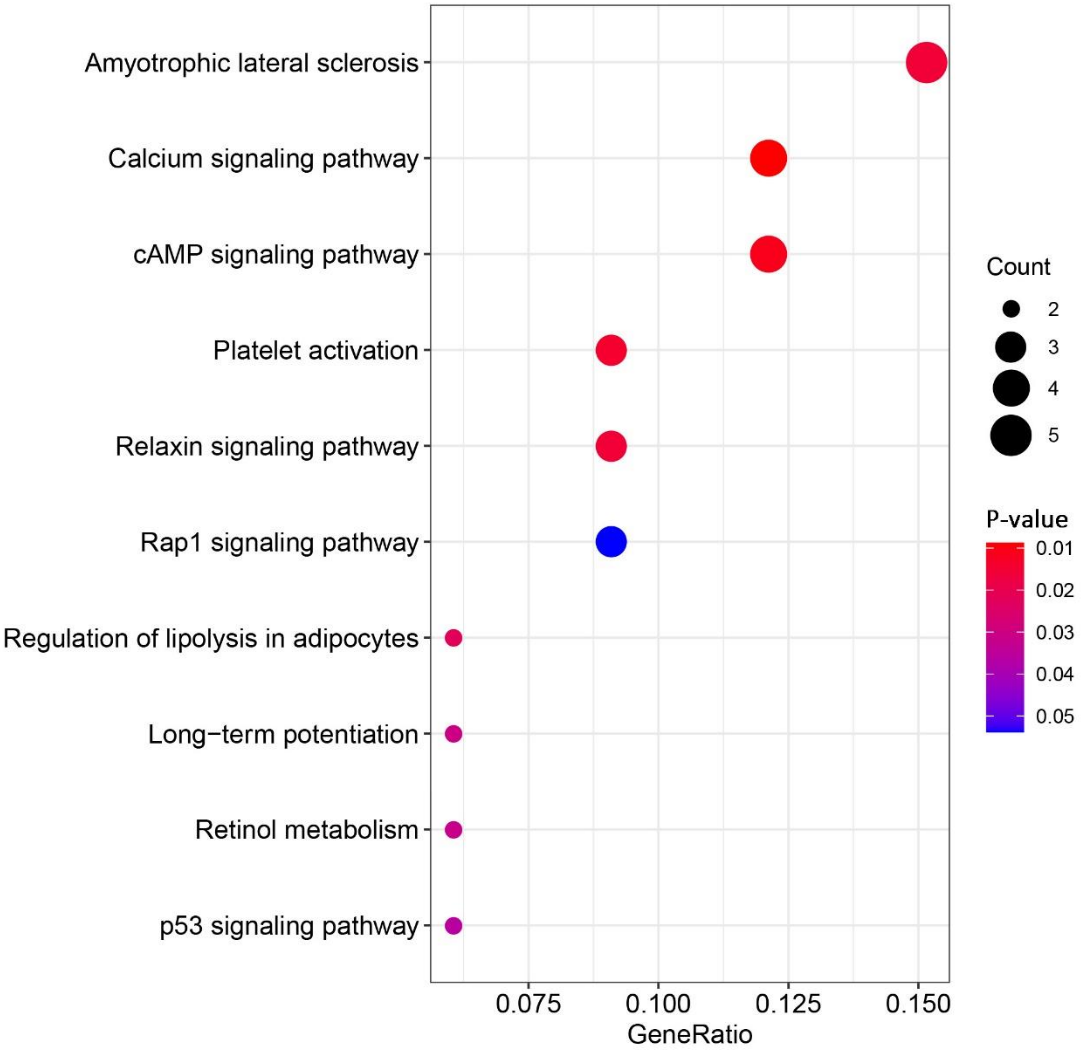


Figure S26. The pathway enrichment analysis plot of the hypermethylated down-regulated genes.

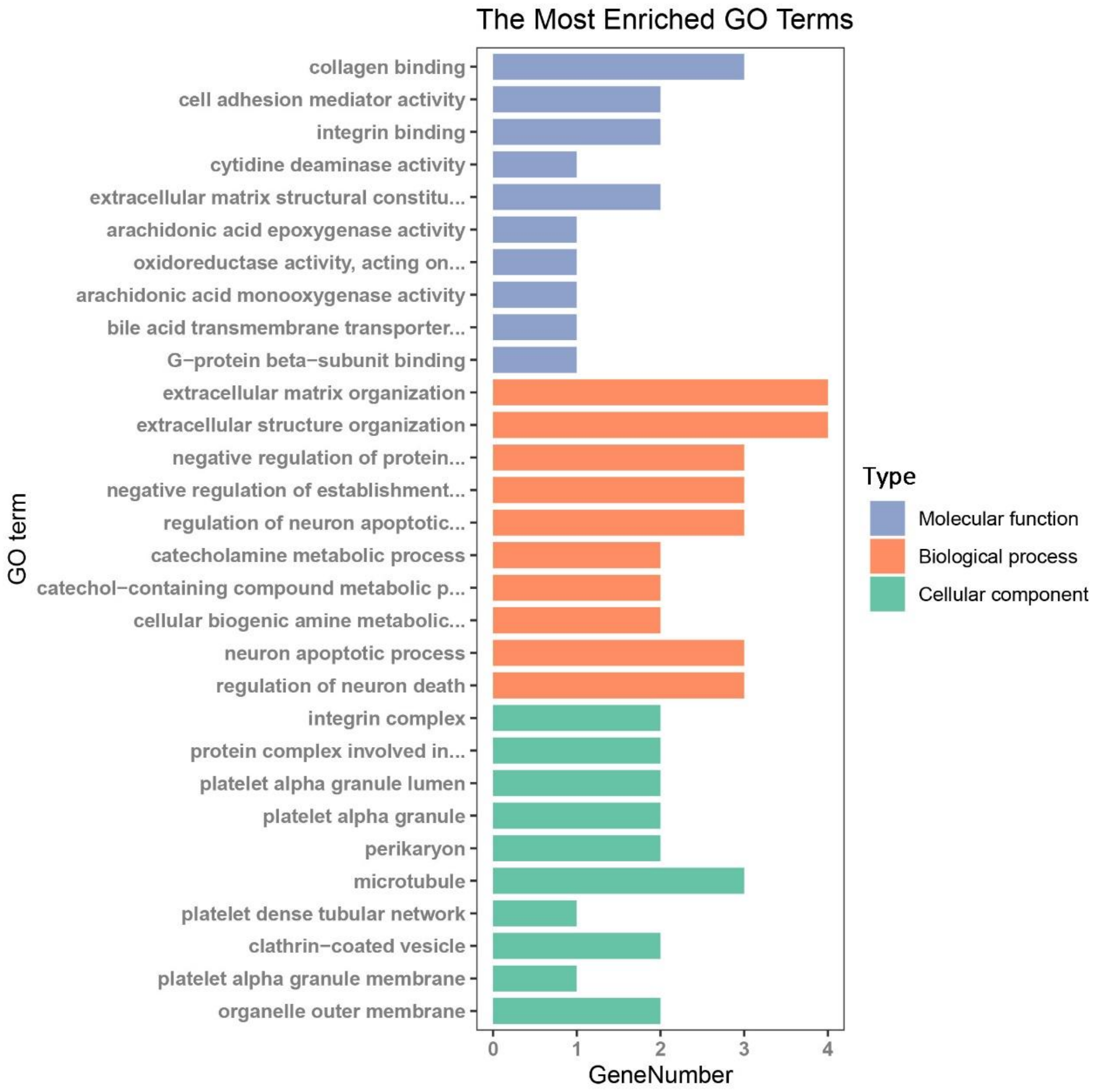


Figure S27. The GO enrichment analysis plot of the hypomethylated up-regulated genes.

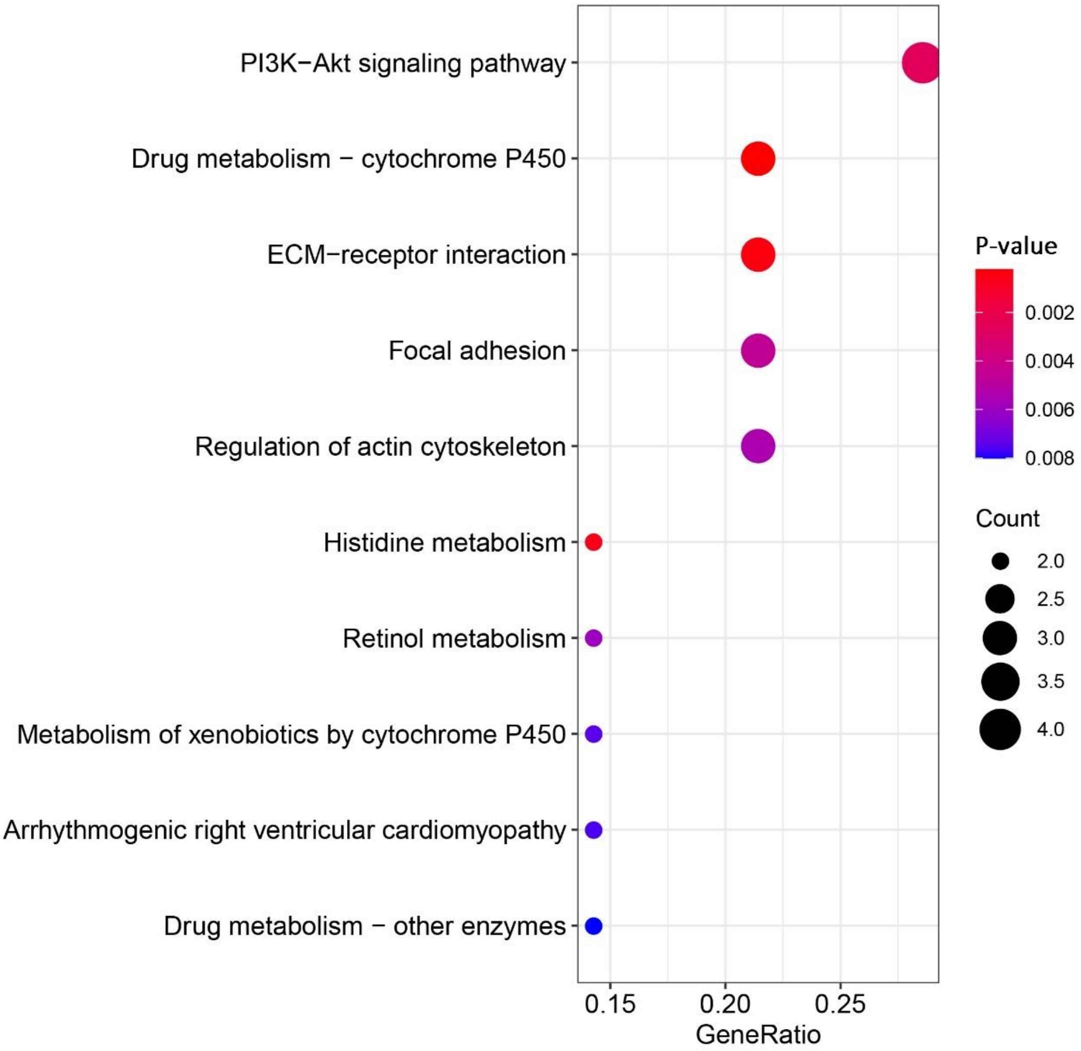


Figure S28. The Pathway enrichment analysis plot of the hypomethylated up-regulated genes.

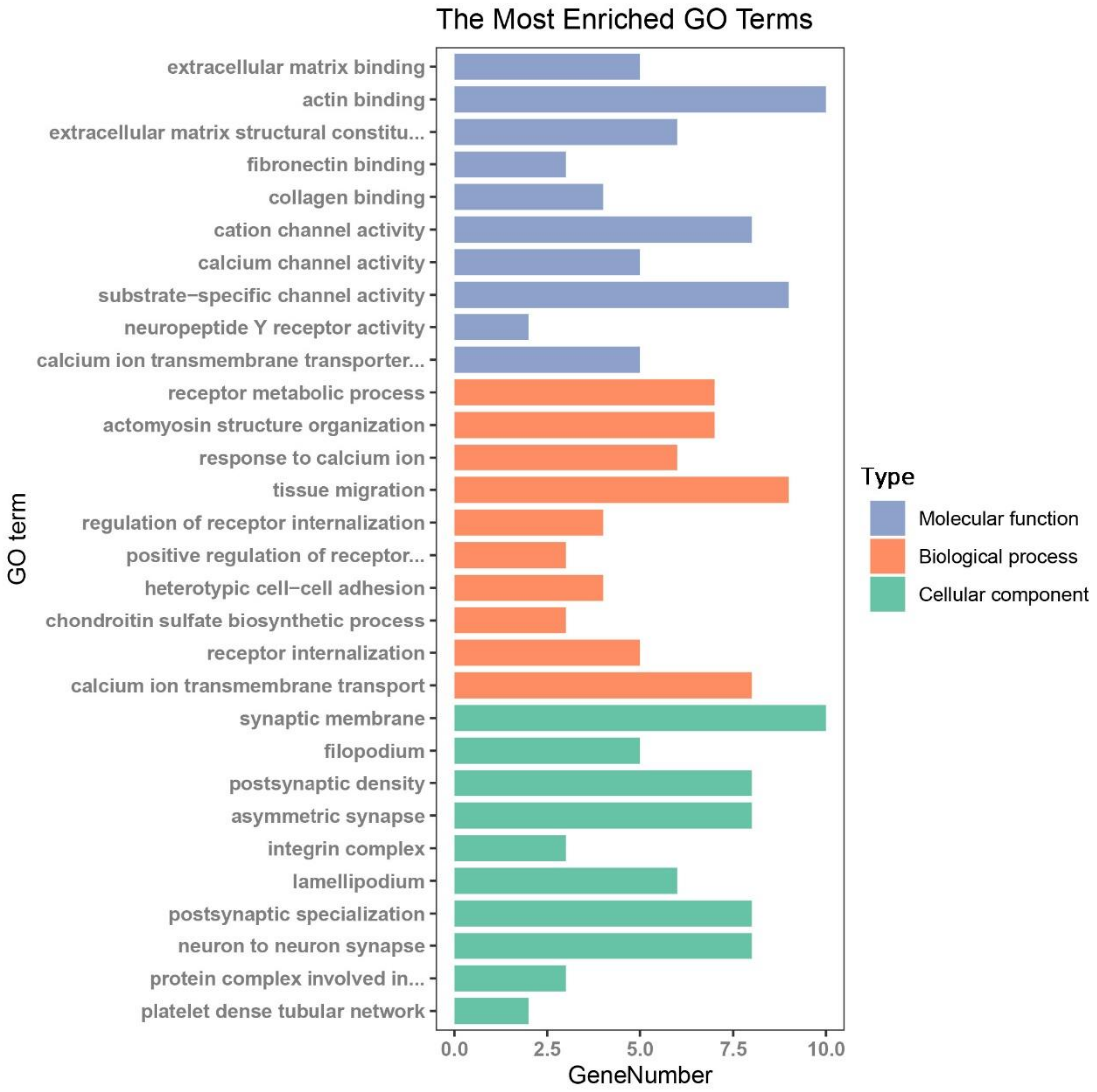


Figure S29. The GO enrichment analysis plot of the overlapped DEGs between EXP-Blood-HIV-Resistance and the ChIP-Seq data.

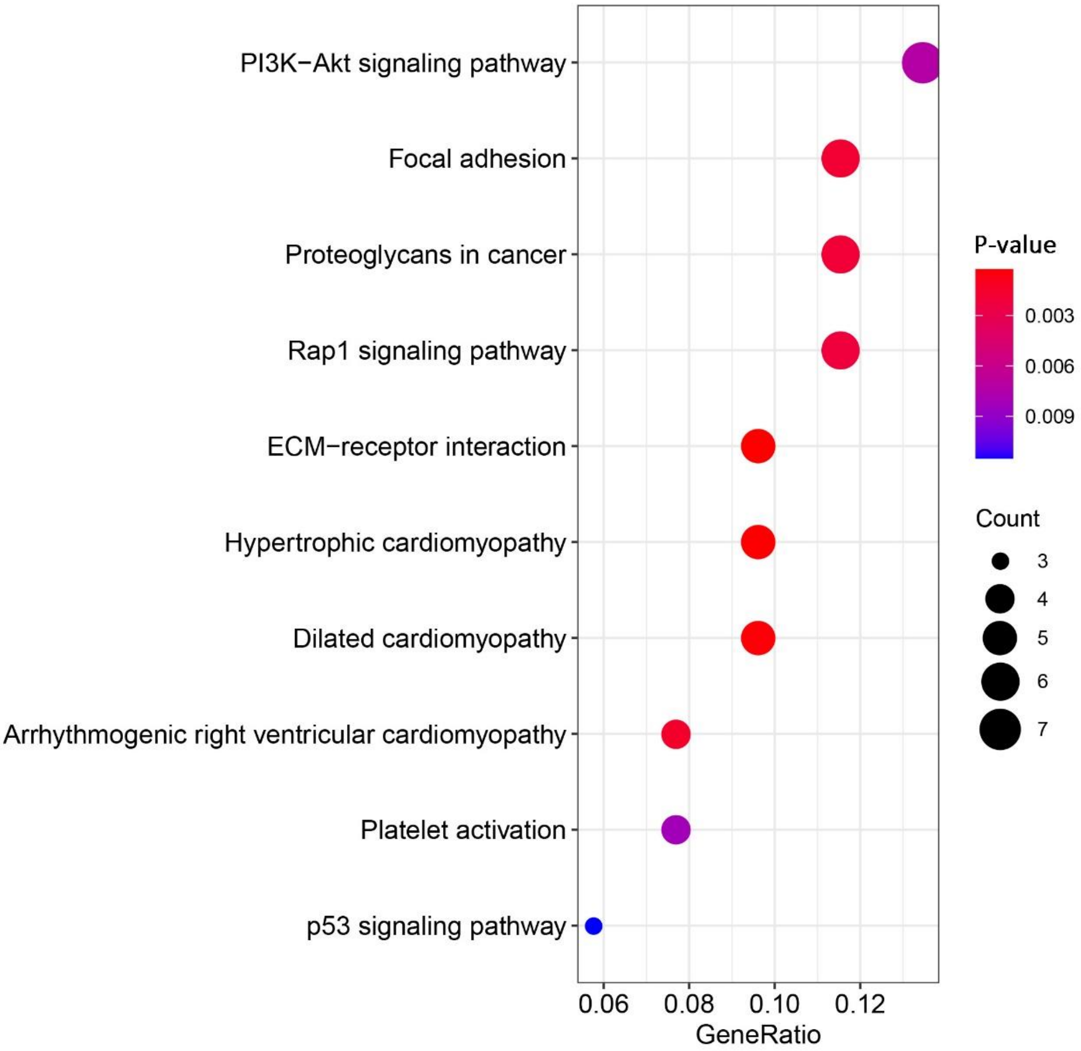


Figure S30. The pathway enrichment analysis plot of the overlapped DEGs between EXP-Blood-HIV-Resistance and the ChIP-Seq data.

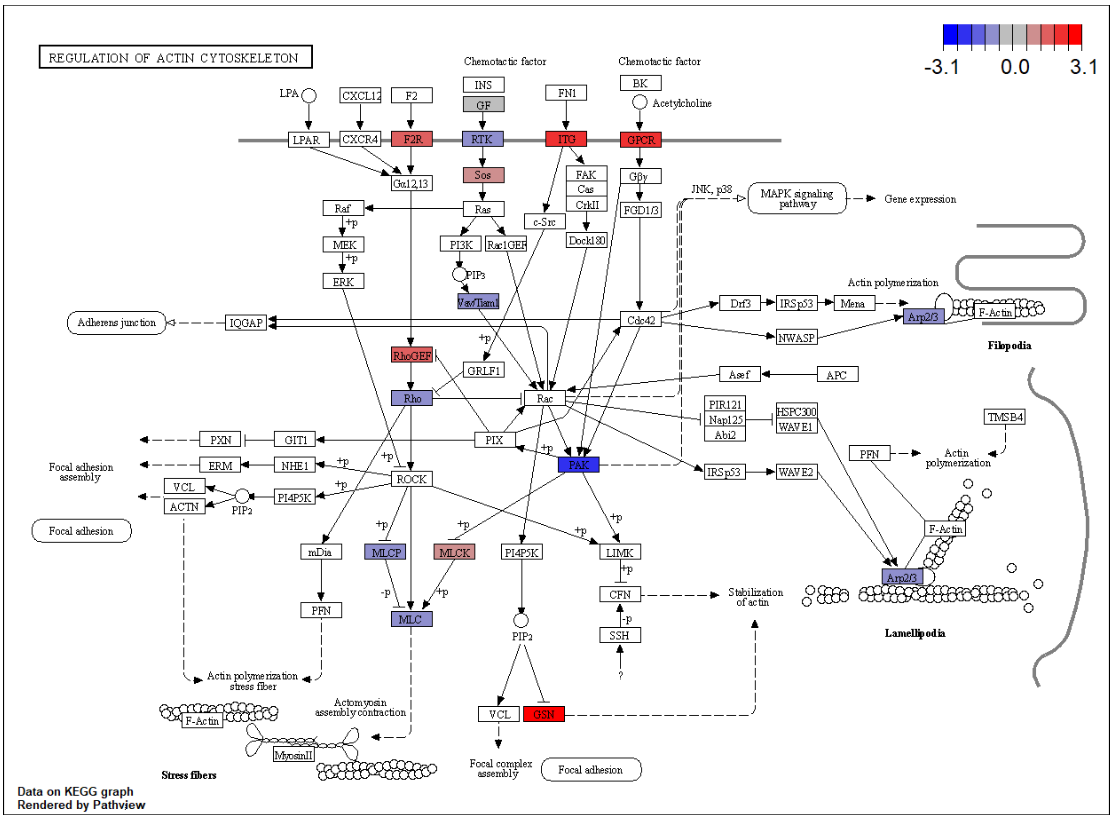


Figure S31. The expression of DEGs in EXP-Blood-HIV-Resistance and EXP-CD4-HIV-Resistance in the pathway, regulation of actin cytoskeleton.

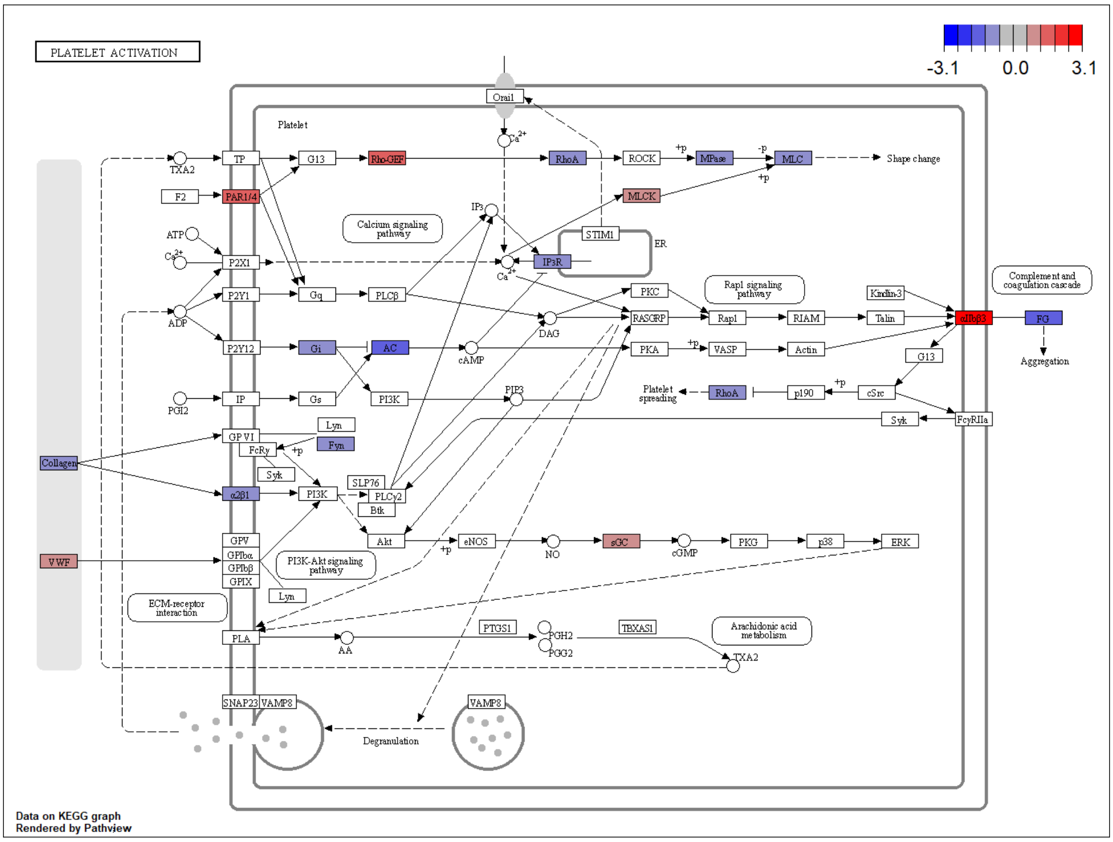


Figure S32. The expression of DEGs in EXP-Blood-HIV-Resistance and EXP-CD4-HIV-Resistance in the pathway, platelet activation.

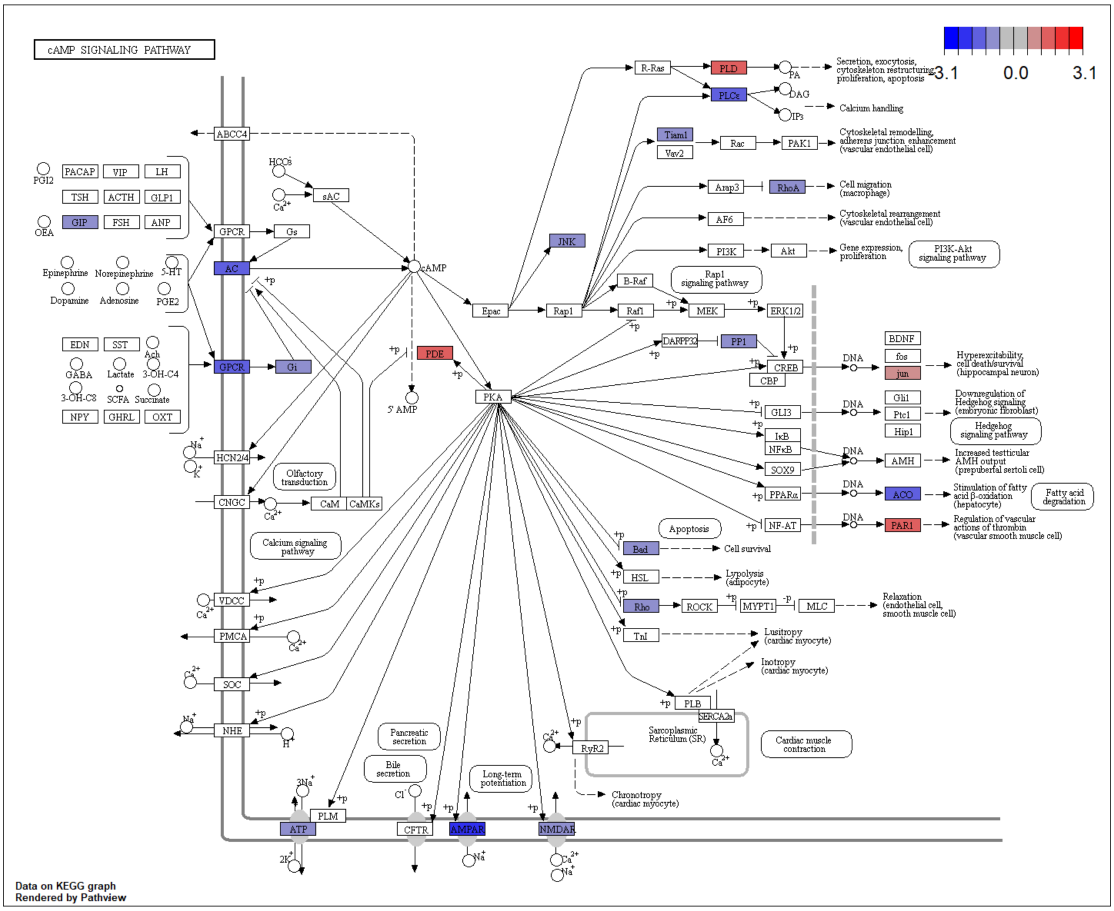


Figure S33. The expression of DEGs in EXP-Blood-HIV-Resistance and EXP-CD4-HIV-Resistance in the pathway, cAMP signaling pathway.

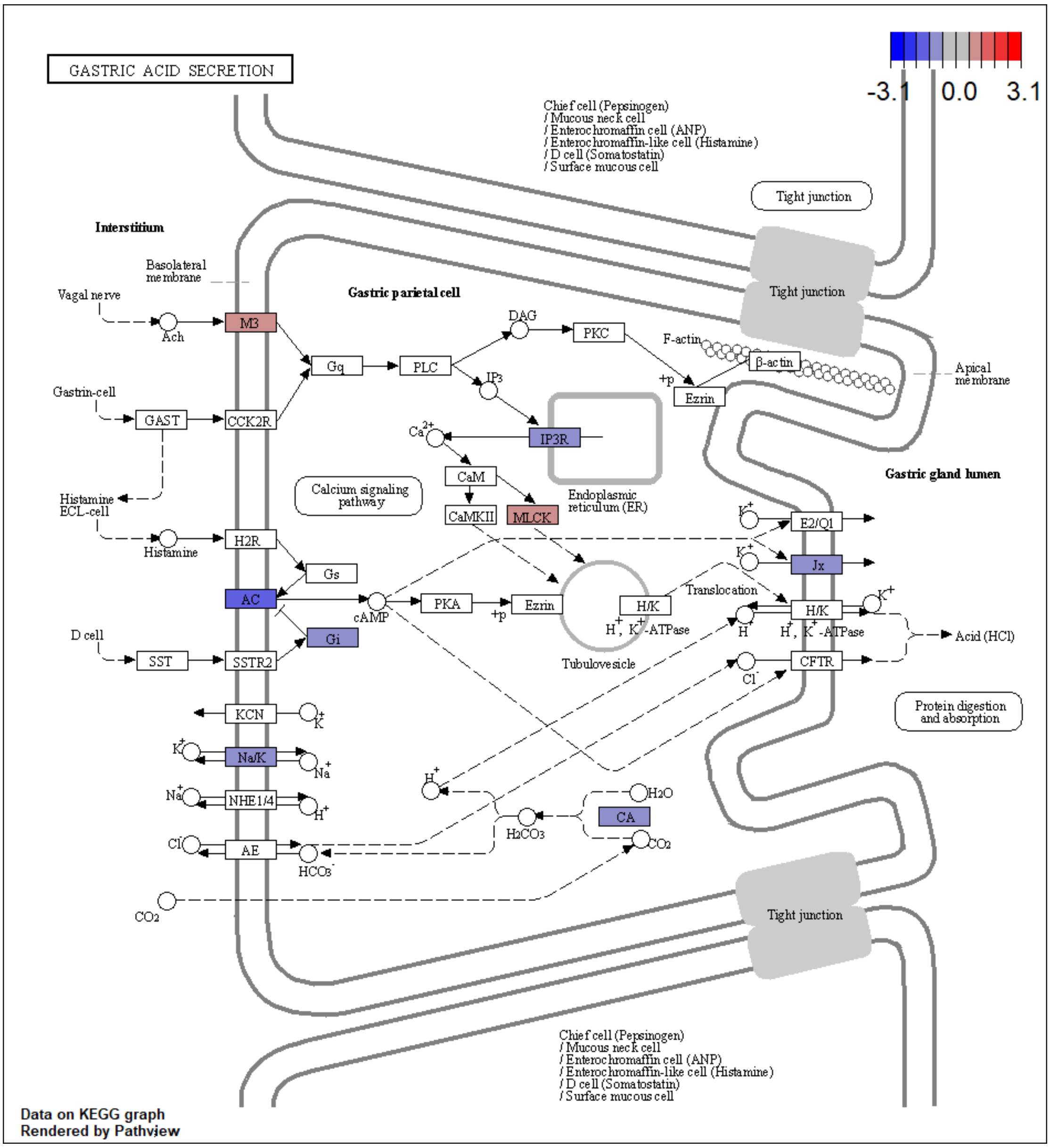


Figure S34. The expression of DEGs in EXP-Blood-HIV-Resistance and EXP-CD4-HIV-Resistance in the pathway, gastric acid secretion.

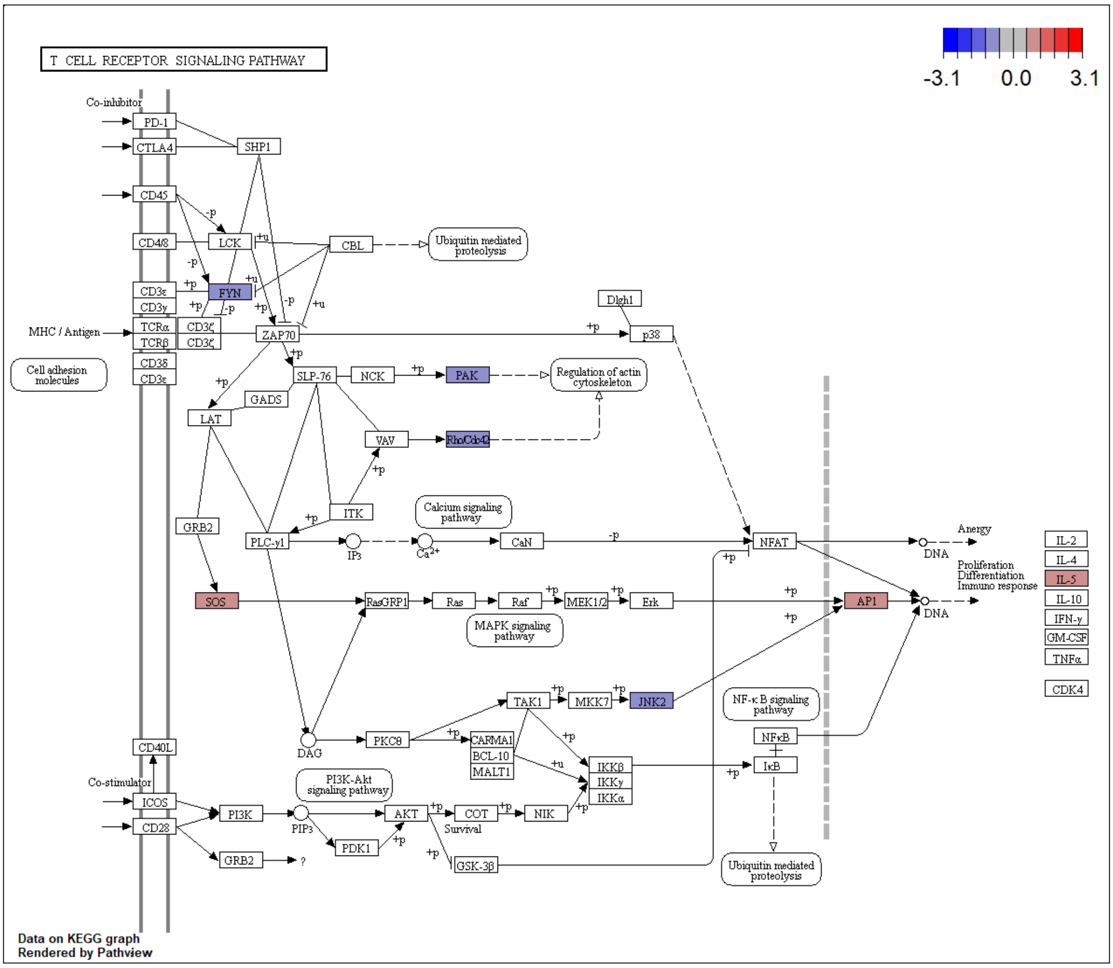


Figure S35. The expression of DEGs in EXP-CD4-HIV-Resistance in TCR Signaling Pathway.
